# Convergent evolution in SARS-CoV-2 Spike creates a variant soup that causes new COVID-19 waves

**DOI:** 10.1101/2022.12.05.518843

**Authors:** Daniele Focosi, Rodrigo Quiroga, Scott A. McConnell, Marc C Johnson, Arturo Casadevall

## Abstract

The first 2 years of the COVID-19 pandemic were mainly characterized by convergent evolution of mutations of SARS-CoV-2 Spike protein at residues K417, L452, E484, N501 and P681 across different variants of concern (Alpha, Beta, Gamma, and Delta). Since Spring 2022 and the third year of the pandemic, with the advent of Omicron and its sublineages, convergent evolution has led to the observation of different lineages acquiring an additional group of mutations at different amino acid residues, namely R346, K444, N450, N460, F486, F490, Q493, and S494. Mutations at these residues have become increasingly prevalent during Summer and Autumn 2022, with combinations showing increased fitness. The most likely reason for this convergence is the selective pressure exerted by previous infection- or vaccine-elicited immunity. Such accelerated evolution has caused failure of all anti-Spike monoclonal antibodies, including bebtelovimab and cilgavimab. While we are learning how fast coronaviruses can mutate and recombine, we should reconsider opportunities for economically sustainable escape-proof combination therapies, and refocus antibody-mediated therapeutic efforts on polyclonal preparations that are less likely to allow for viral immune escape.

## Introduction

In the third year of the COVID-19 pandemic, the majority of the general population is now largely protected from severe COVID-19 disease and death by mass vaccination campaigns and by immunity from former infection. Unfortunately, SARS-CoV-2 remains a life-threatening pathogen for immunocompromised (IC) patients who are unable to mount a protective immune response. IC individuals create a cohort population in whom the virus can persistently replicate, which is a novelty for pandemics. In this regard, advancements in therapeutics and supportive care have greatly increased the prevalence of IC patients compared to just a few decades ago. SARS-CoV-2 infection in IC patients is arguably the most difficult current problem in the COVID-19 pandemic for these individuals can have large viral loads with inevitably include antigenically different viruses and have a diminished capacity for clearing the infection.

Since Summer 2022, SARS-CoV-2 transmission has proceeded undisturbed worldwide after the relaxation of nonpharmaceutical interventions such as lockdowns, social distancing, hand hygiene, and face masks, which together with the waning of infection- and vaccine-elicited immunity, has increased opportunities for spread and the number of susceptible individuals, respectively. Hence, the increase in the “human culture medium” has led to large infectious waves during 2022, with estimated excess deaths similar to those observed in 2020 [1]. While acquisition and waning of immunity from former infections is not a novel occurrenece, the COVID-19 pandemic has created conditions whereby the natural course of a coronavirus pandemic is changed by introducing timely vaccination campaigns and therapeutics targeting the viral receptor domain. There is no historical precedent for the current situation. The combined action of increasing cumulative viral loads in the “human culture medium” and such selective pressures has led to an unprecedented increase in viral diversification in 2022. WHO nomenclature for variants of concern remained stuck at “Omicron”[2], while alternative naming schemes introduced novel names to designate lineages that are responsible for thousands of hospitalizations. The most refined phylogeny to date has been released by PANGOLIN which counts more than 600 designated Omicron sublineages at the time of writing (https://www.pango.network/summary-of-designated-omicron-lineages/), accounting for more than 45% of SARS-CoV-2 variability (Figure 1). Of interest, such increase in divergence was detected despite a 75% reduction in genomic surveillance in 2022, which is mainly due to budget constraints. After peaking at 1 million sequences in January 2022, the number of new sequences deposited at the site decreased to t 250,000 in October 2022 (https://cov-spectrum.org/explore/World/AllSamples/Past6M/sequencing-coverage). Consequently, it is likely that the number of defined circulating sublineages is an underestimate of the viral genetic variation in the current pandemic.

**Figure 1.**
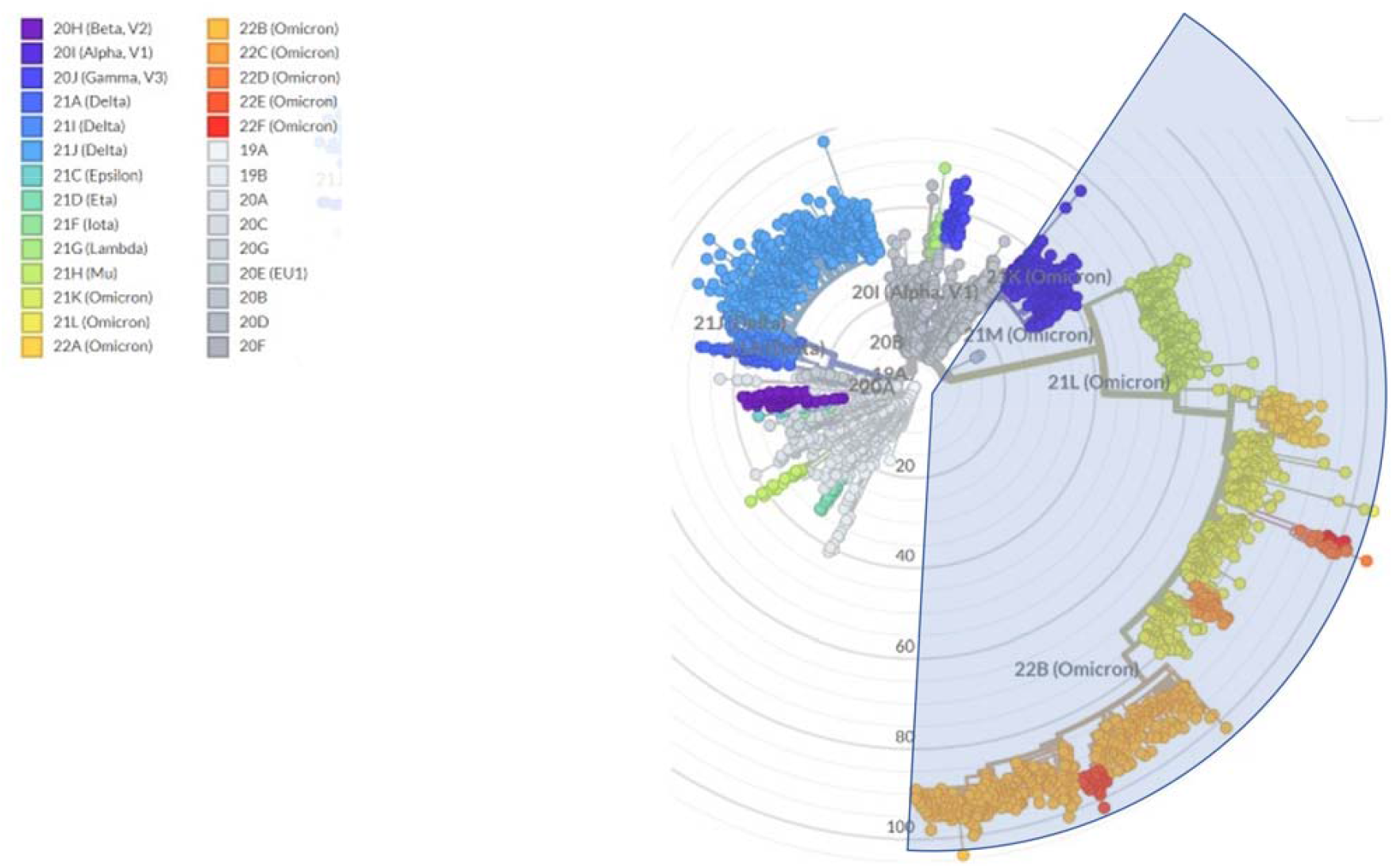
Radial tree of SARS-CoV-2 evolution, with branch length approximating divergence, showing that Omicron (light blue shadow) currently includes more than 45% or variations across 3045 genomes sampled between Dec 2019 and Nov 2022. Accessed online at https://nextstrain.org/ncov/gisaid/global/all-time?l=radial&m=div on November 26, 2022.

### Mutation rates and mutational spectra

Mutation rate (MR) is often used interchangeably to indicate 2 different things: occurrence of mutations within a single host (intrahost evolution at individual level without any demand for outcompeting co-circulating strains) or step-wise accumulation of mutations (“antigenic drift”) that get fixated within a species. While the first meaning has been demonstrated (e.g., in IC hosts[3-5], and after administration of the small molecule antiviral molnupiravir which known to increase G→A and C→U transition mutations[6], potentially contributing to new linages), from an evolutionary standpoint it is the second meaning which is more interesting and already well-established for other respiratory pathogens[7], including the related human coronavirus 229E[8].

Early in the pandemic, data suggested that mass vaccination could restrict SARS-CoV-2 mutation rates (MR): the diversity of the SARS-CoV-2 lineages declined at the country-level with increased rate of mass vaccination (*r* = -0.72) and vaccine breakthrough patients harbor viruses with 2.3-fold lower diversity in known B cell epitopes compared to unvaccinated COVID-19 patients [9]. Also, vaccination coverage rate was inversely correlated to the MR of the SARS-CoV-2 Delta variant in 16 countries (*r*^*2*^= 0.878) [10].

Ruis *et al* found a halving in the relative rate of G→T mutations in Omicron compared to pre-Omicron sublineages[11]. To exclude selective pressures on the derived protein structures, Bloom *et al* found similar results by repeating the analysis focusing on 4-fold degenerate codons (i.e. codons that can tolerate any point mutation at the third position, although codon usage bias restricts this in practice in many organisms) [12]. Replicaion of viruses and bacteria in the lower respiratory tract has been associated with high levels of G>T mutations and for SARS-CoV-2 this effect occurred with Delta but was lost in Omicron [11]. Such changes on mutation type and rate could theoretically stem from from mutations affecting genome replication and packaging [13], as well as from mutations in genes encoding proteins (e.g. APOBEC) that antagonize host innate-immune factors, which otherwise will mutate viral nucleic acids[14-16] and/or from environmental factors [6].

The average MR of the entire SARS-CoV-2 genome was estimated from the related mouse hepatitis virus (MHV) to be 10^−6^ nucleotides per cycle, or 4.83 × 10^−4^ subs/site/year, which is similar, or slightly lower, that observed for other RNA viruses [17]. Following the removal of mandatory nonpharmaceutical interventions such as face masks, social distancing, and quarantine in most western countries, vaccination was not sufficient to prevent hyperendemicity. The MR of SARS-CoV-2 consequently doubled from 23 substitutions per year before December 2021 to 45 substitutions per year after December 2021, coinciding with the advent of omicron (Figure 2), which approximates 14.5/subs/year for the ∼30 kb SARS-CoV-2 genome. This rate should set the upper limit for mutation frequency, as many mutations will not be viable and/or transmissible, and thus not observed in the sequencing data at baseline. It had been previously shown that the P203L mutation in the error-correcting exonuclease non-structural protein 14 (nsp14) almost doubles the genomic MR (from 20 to 36 SNPs/year) [18]. While this change is not prevalent in Omicron lineages, many changes in the replication machinery appeared with Omicron, such as K38R, Δ1265, and A1892T in Nsp3; P132H in Nsp5; I189V in Nsp6; P323L in Nsp12; and I42V in Nsp14, and some of them could have contributed to the MR jump[19].

**Figure 2.**
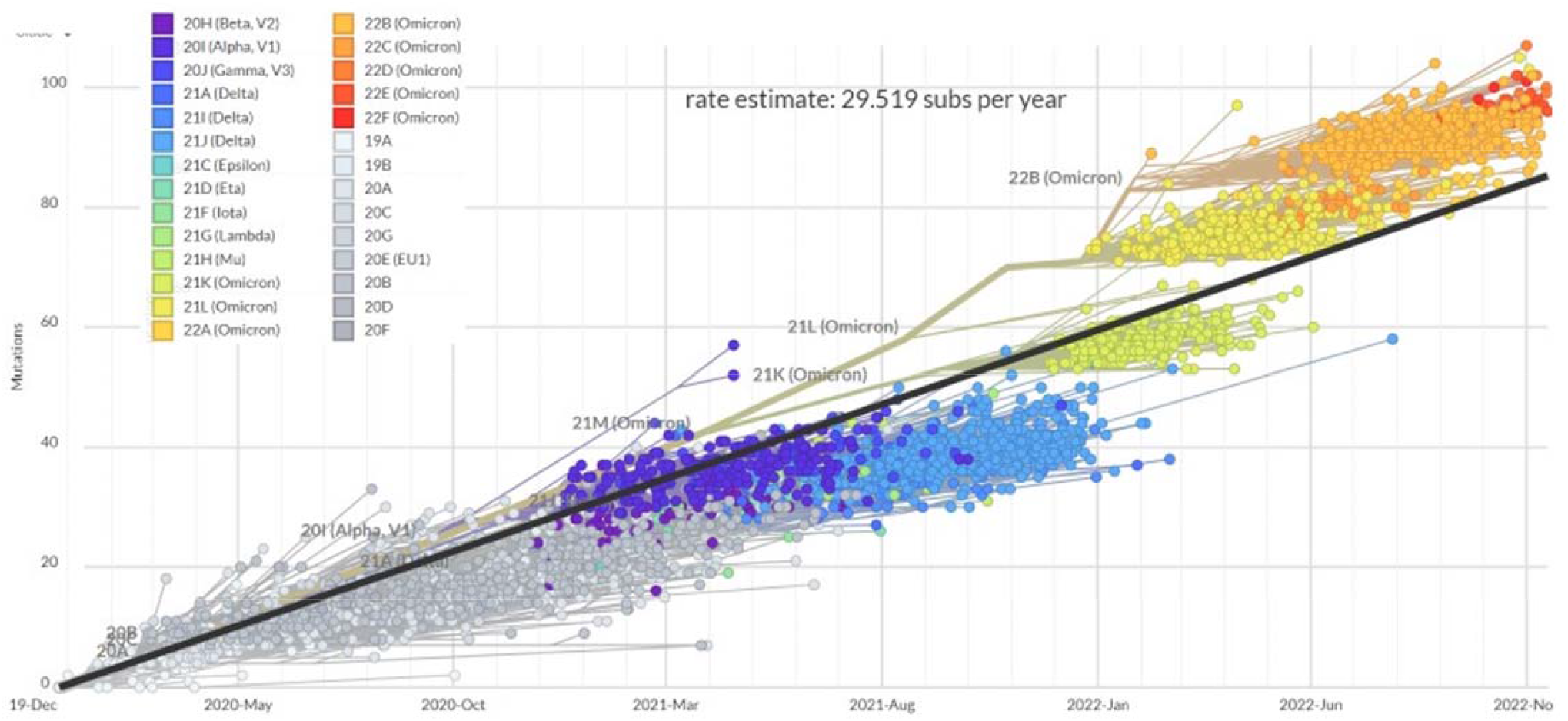
Clock tree of SARS-CoV-2 evolution, with regression line showing an increase in the estimate rate of substitutions per year across 3045 genomes sampled between Dec 2019 and Nov 2022. Accessed online at https://nextstrain.org/ncov/gisaid/global/all-time?l=clock&m=div on November 26, 2022.

### Convergent evolution

In the midst of such massive lineage divergence, convergent evolution towards certain motifs has become increasingly manifest.

In the pre-Omicron and pre-vaccine era, variants of concern (VOCs) notably converged to mutations which resulted in the following amino acid changes: K417N, L452R, E484K, N501Y, and P681X[20]. These amino acid changes have been proposed to increase the stability of the trimeric protein[21-23], and they emerged in the absence of significant selective pressures by the immune system. K417N, E484A, N501Y and P681H remained hallmarks of BA.2.*, while the paraphyletic BA.4/5 acquired L452R and F486V and the Q493R reversion.

In the last year the BA.2 variant first generated a wave that led first to the paraphyletic BA.4/5 sublineage, which was later joined by a return of so-called “second-generation” BA.2 sublineages (Figure 3), with BA.2.75.* and BA.2.3.20 being the most circulated. Since Summer 2022, each of those sublineages has amazingly converged with changes at the receptor-binding domain (RBD) residues R346, K444, L452, N450, N460, F486, F490, Q493, and S494 (see Supplementary Table 1)[24]. E484A remained instead stable, with 484K never detected, A484G seen only in BA.2.3.20, and A484T seen only in XBB.1.3. More recently, convergence in indels within the N-terminal domain (NTD), as previously recognized in Brazilian VOCs[25], was reported for Omicron sublineages: in particular, Y144del has been found in BA.4.6.3, BJ.1, BU.1, BQ.1.8.*, BQ.1.1.10, BQ.1.1.20, BQ.1.18, and XBB.*)[26].

**Figure 3.**
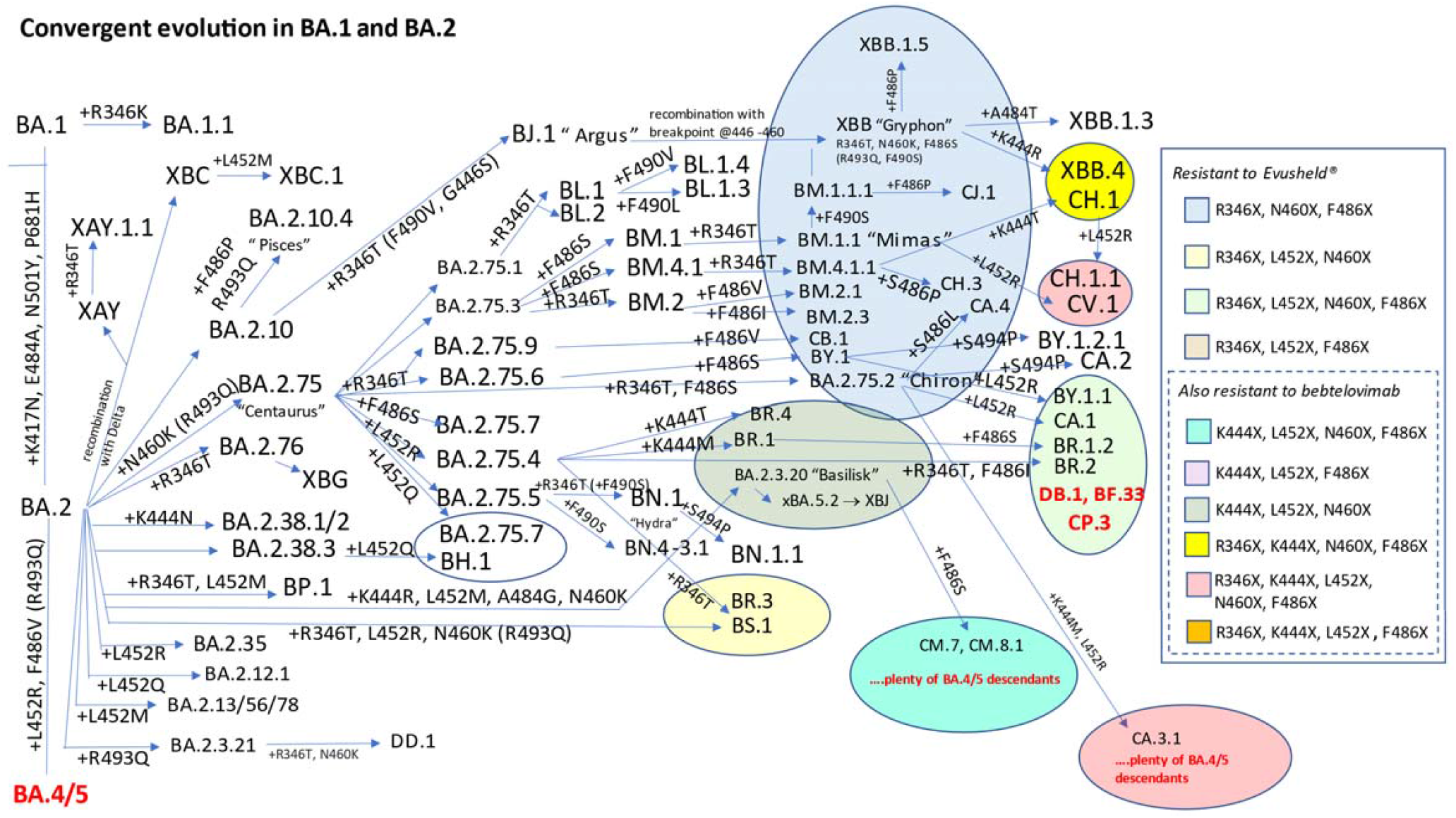
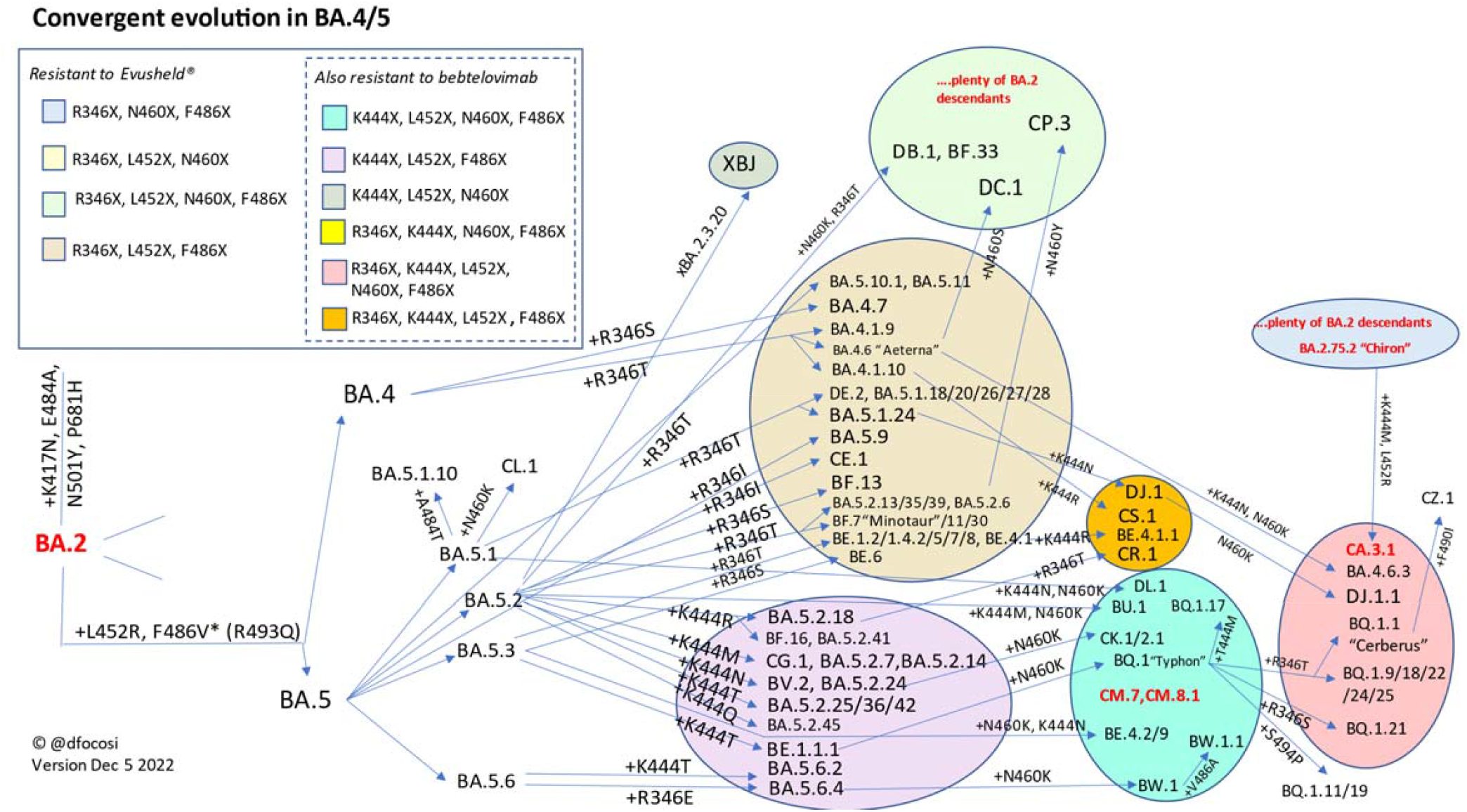
Diagram representing all SARS-CoV-2 Omicron sublineages designated by PANGOLIN as of November 26, 2022 for which at least one of the Spike RBD immune escaping mutations (R346X, K444X, L452X, N460X, F486X, or R493Q) represents a branching event. Mythological names introduced by Ryan T Gregory and used colloquially are also reported. Convergence towards combos of this mutations is noted, with different background colors representing different combinations. Resistance of each combination to clinically authorized anti-Spike mAbs is reported on the right box. For visualization purposes, the upper panel shows BA.1 and BA.2 evolution, while the lower panel shows BA.4/5 evolution.

This “variant soup” can be organized and stratified according to the number of key Spike mutations present, and although the number of key mutations acquired correlates well with increasing fitness, this is only so within each lineage, which shows that the biology of SARS-COV-2 infection goes beyond what occurs in the Spike protein (Figure 4). At present, only the BQ.1-derived lineages with 7 or more selected mutations display a clear relative growth advantage relative comparison to the BQ.1.1 baseline. Convergence was clearly observed at the amino acid level, with different nucleotide mutations leading to similar amino acid changes: e.g., N460K was caused by T22942A in BQ.1*, XAW and some of the BA.5.2 sublineages, while it was caused by T22942G in BA.2.75*(all lineages), BA.2.3.20, BS.1, BU.1, XBB, XAK and BW.1 (BA.5.6.2.1). Another impressive example of this convergent evolution is the <u>S</u>pike of BA.4.6.3, BQ.1.18 and BQ.1.1.20 independently acquiring the following amino acid changes since their last shared common ancestor: Y144del, R346T, N460K, L452R, F486V and the R493Q reversion. Also, BA.4.6.3 has acquired K444N, while BQ.1.18 and BQ.1.1.20 acquired K444T.

**Figure 4.**
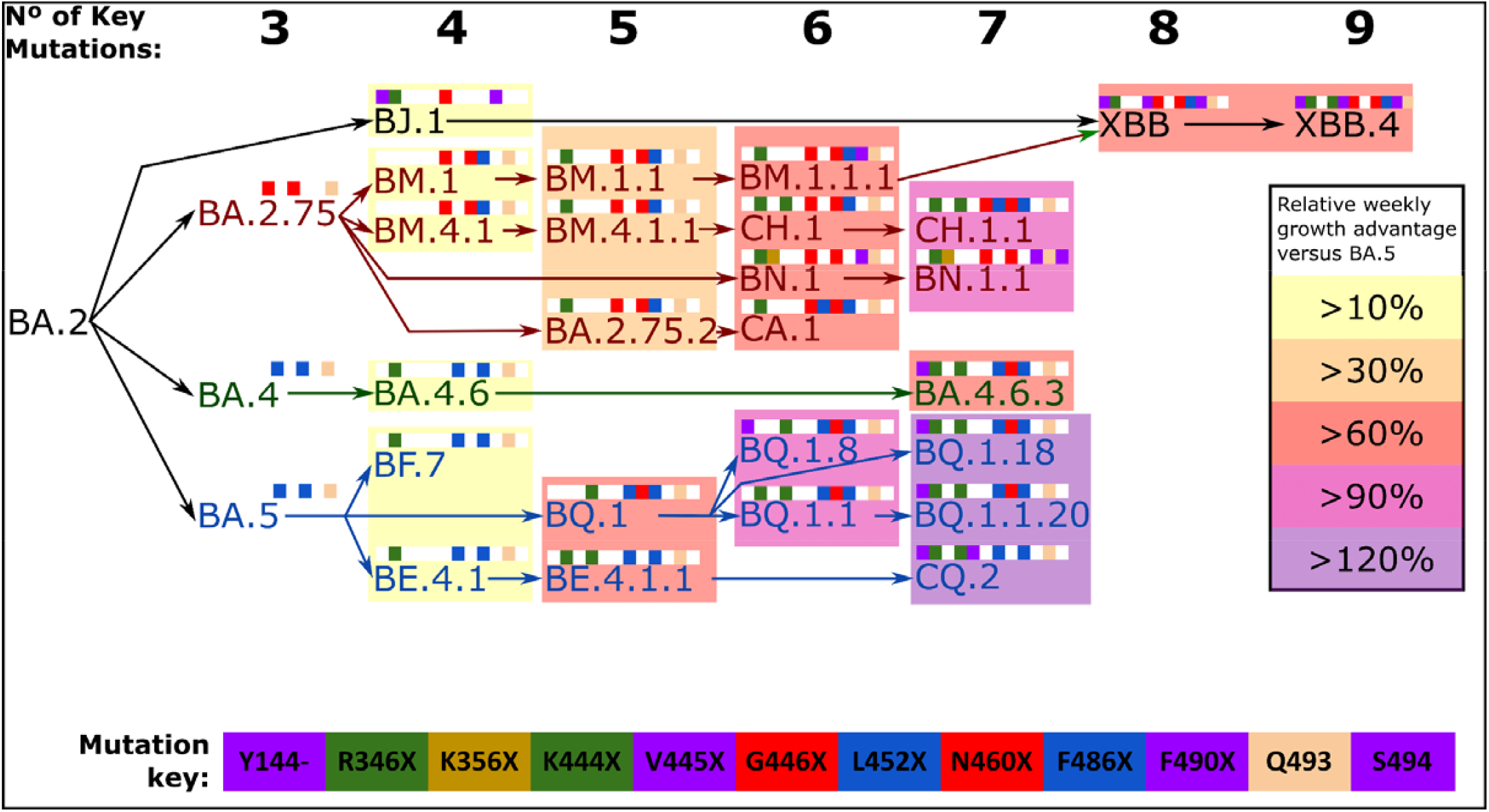
Step-wise accumulation of key Spike mutations involved in immune escape within SARS-CoV-2 Omicron sublineages increase the relative rowth rate. Lineage name text is color coded, where BA.5 descendants are in blue text, BA.4 descendants in green text and BA.2.75 descendants are in r d text. Each mutation is color coded as shown in the mutation key, and depicted as colored squares when present or white squares if absent. Number of k y mutations of each lineage is summarized at the top. Relative growth rates were calculated using BA.5 lineage as baseline, for groups of BA.4, BA.5, BA.2.75 and XBB descendant lineages with each exact total number of key mutations. Relative growth rates were calculated using CoV-Spectrum [67]

#### Escalating immune escape

SARS-CoV-2 evolution represents an accelerated movie of Darwinian selection. Variants that are more likely to escape vaccine- and infection-elicited immunity that are more fit expand at the expense of those less fit. While it may sound obvious, we now have formal evidence of such evolution, with PANGOLIN descendants invariably having increased RBD immune escape scores compared to parental strains (**Figure 5**). In this ongoing race, descendants invariably replace parents, as these are fitter in hosts with pre-existing immunity. In this regard, the chances for saltations lineages that emerged after intrahost evolution in IC patients (i.e. in the absence of RBD immune escape) seem minimal: accordingly, despite the initial hypothesis of intrahost evolution to explain the saltation seen with the emergence of Omicron, recent evidence suggests that Omicron ancestors circulated undetected long before the exponential spread [27].

**Figure 5.**
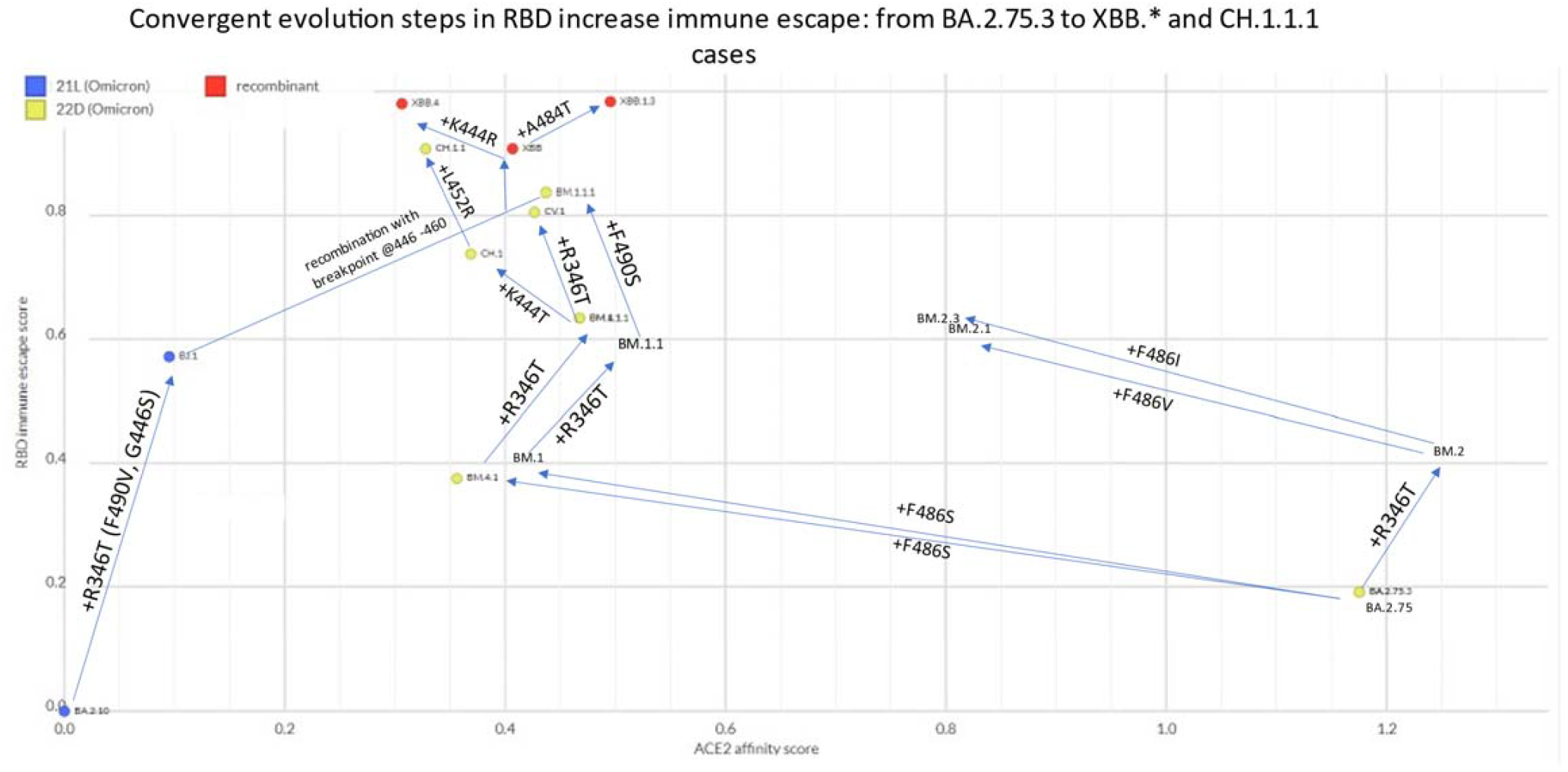
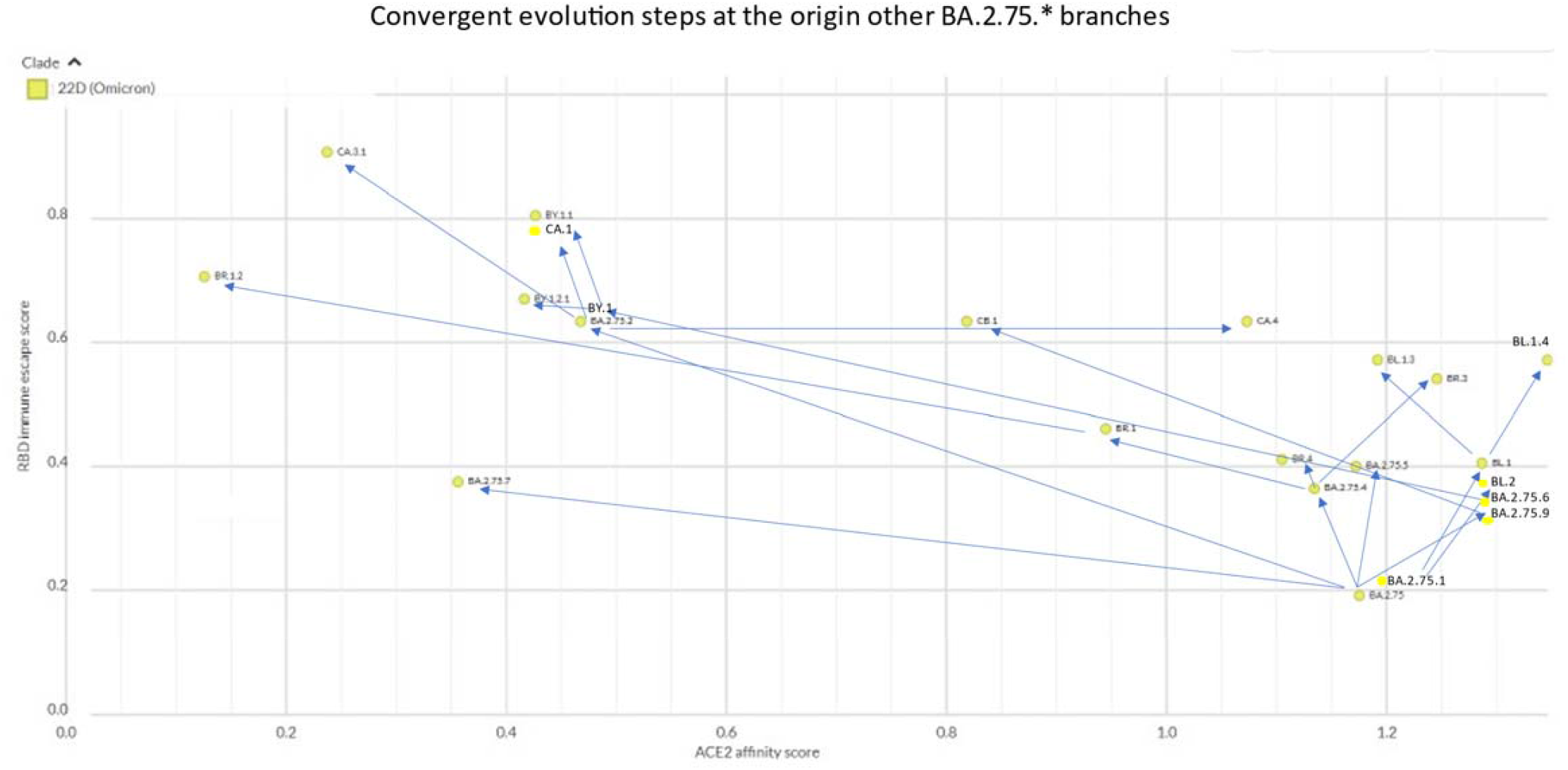
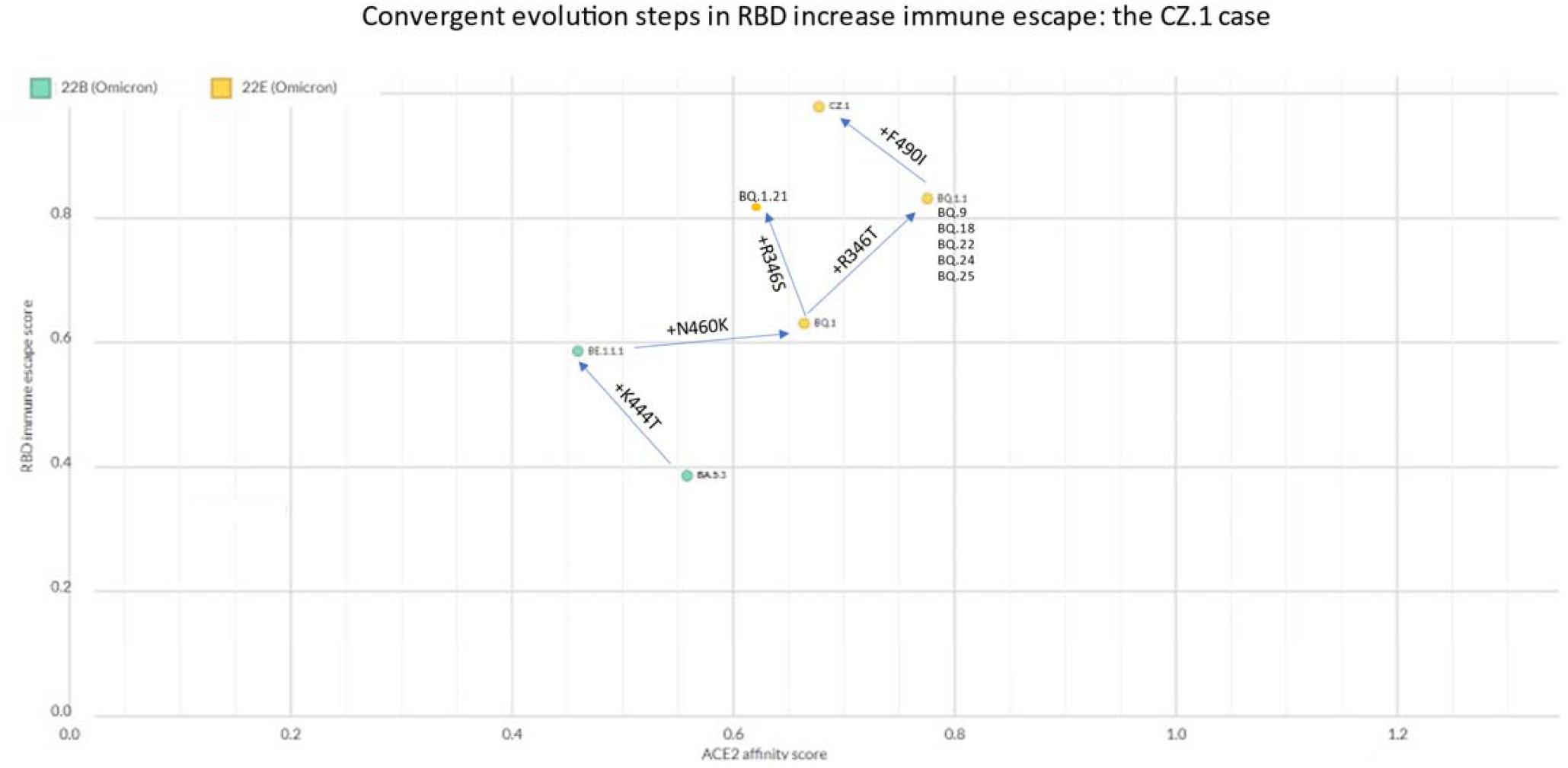
Evolutionary steps at the basis of the major Omicron branches. (CZ.1, XBB.* and CH.1.1.1, and other BA.2.75.* descendants), showing progressive increases in RBD immune escape score (as calculated here: https://jbloomlab.github.io/SARS2_RBD_Ab_escape_maps/escape-calc/). Chart created on NextStrain [68] (https://next.nextstrain.org/staging/nextclade/sars-cov-2/21L?gmin=15&l=scatter&scatterX=ace2_binding&scatterY=immune_escape&showBranchLabels=all)

RBD immune escape can nowadays be estimated *in silico* based on *in vitro* data (https://jbloomlab.github.io/SARS2_RBD_Ab_escape_maps/escape-calc/). RBD immune escape is clearly a moving scale with an evolving asymptote. E.g., by changing vaccine composition [28] we are likely to reset the “game”.

### ACE2 affinity fine tuning

ACE2 affinity can be estimated *in silico* (https://github.com/jbloomlab/SARS-CoV-2-RBD_DMS_Omicron/blob/main/results/final_variant_scores/final_variant_scores.csv). Several Omicron sublineages showed remarkable examples of further evolution at Spike residues that were already recently mutated. E.g.,

- BQ.1 already had K444T inherited from BE.1.1.1, but further mutated into 444M in the child BQ.1.1.17
- XBB.1 already had E484A inherited from the BA.2 parent, but further mutated into 484T in the child XBB.1.3
- BA.2.3 already had E484A inhertited from the BA.2 parent, but further mutated into 484G in the child BA.2.3.20, which caused an impressive increase in ACE2 affinity (to whom K444R, L452M, and N460K contributed)
- BM.4.1.1 already had F486S inherited from the BM.4.1 parent but further mutated into 486P in CH.3
- BM.1.1.1 already had F486S inherited from the BM.1 parent but further mutated into 486P in the child CJ.1
- XBB.1 already had F486S inherited from the BM.1.1.1 parent but further mutated into 486P in the child XBB.1.5
- BA.2.75.2 already had F486S inherited from the BA.2.75 parent, but further mutated into 486L in the child CA.4
- BA.5.2.1 already had F486V inherited since BA.5, but further mutated to 486I in BF.12
- BW.1 already had F486V inherited from the BA.5 parent, but further mutated into 486S in the child BW.1.1

Seven of these examples manifest escalating affinities for ACE2, with the other 2 representing no change in ACE2 affinity (**Figure 6**).

**Figure 6.**
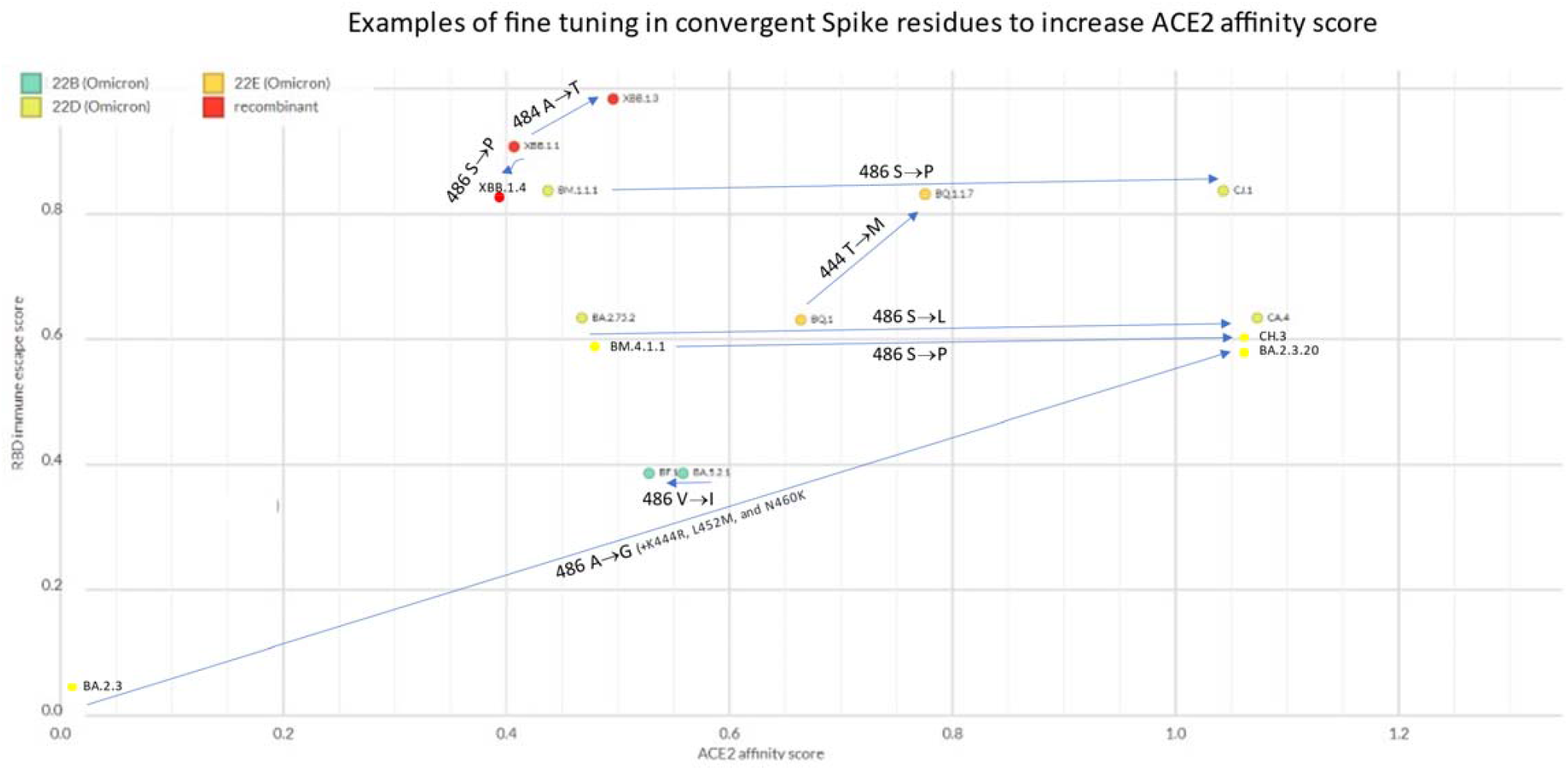
Sequential mutational events at the same Spike amino acid residues showing no change or progressive increases in ACE2 affinity score (as calculated here: https://github.com/jbloomlab/SARS-CoV-2-RBD_DMS_Omicron/blob/main/results/final_variant_scores/final_variant_scores.csv). Chart created on NextStrain [68] (https://next.nextstrain.org/staging/nextclade/sars-cov-2/21L?gmin=15&l=scatter&scatterX=ace2_binding&scatterY=immune_escape&showBranchLabels=all)

### Mutually exclusive mutations

Mutually exclusive mutations across the entire SARS-CoV-2 genome have been previously studied[29], but the vast constellation of Omicron sublineages provides an unique opportunity for an in-depth exploration of substitutions that are incompatible in combination. The best examples so far are N450X and R346X mutations, which have not yet been observed together in more than 6 millions of Omicron sequences. Two dipolar interactions exist between the carboxamide group of Asn and the guanidino group of Arg in the ancestral sequence, stabilizing the receptor binding module (RBM) tertiary fold (Figure 7, left). R346 resides within a short loop between helix α1 and beta strand 1. N450 is a constituent of the extended RBM insertion into the overall five-stranded antiparallel beta-sheet fold of the domain. As the RBM is the critical determinant for the interaction with ACE2, maintaining its optimal conformation through this stabilizing bond is likely to be essential for pathogenesis. N450D is a common substitution among Omicron lineages. This mutation would result in a similarly sized sidechain but different electrostatic properties (carboxamine → carboxylic acid). This substitution would likely result in a stronger interaction with position 450, as one H-bonding is maintained, and one is replaced with ionic salt bridge between the deprotonated oxygen and the basic guanidino group, provided that the residue at position 346 remains Arg. On the other hand, any substitution at position 346, with the exception of Lys, would result in a significantly shorter, non-cationic sidechain, which would abrogate this RBM-stabilizing interaction. R346K would partially maintain this interaction, replacing a bidentate linkage to N450 with a monodentate dipolar interaction. Thus, the observed mutual exclusivity of mutations at these two sites can be rationalized by their contributions to this stabilizing intradomain interaction.

**Figure 7.**
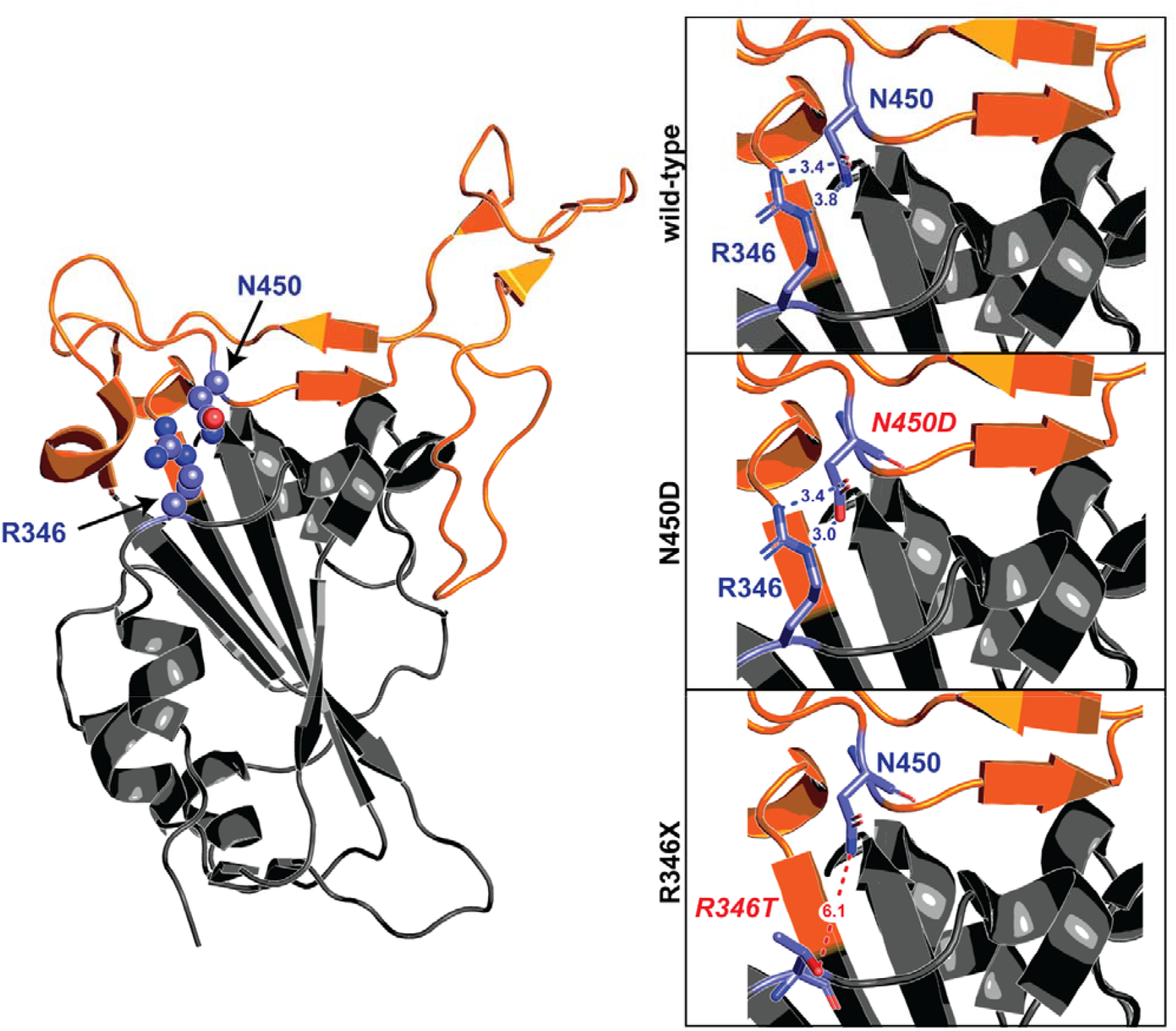
Mutually exclusive mutations at R346 and N450. The receptor binding domain of S is depicted in grey cartoon representation, with the receptor binding module (ACE2 interaction interface) highlighted in orange. Amino acids at the 346 and 450 positions are displayed as purple sticks. A zoomed-in view of the R346-N450 interaction in the ancestral domain, as well as the computationally modelled amino acid substitutions at those two positions, are portrayed in boxes to the right. In the wild-type sequence, the basic R346 sidechain interacts with the N450 residue through a pair of hydrogen bond interactions. N450D results in a similarly sized sidechain, but altered electrostatics. One hydrogen bond is maintained between the neutral oxygen of Asp and Nε of Arg, and a new salt bridge is formed between the anionic deprotonated oxygen of Asp and the cationic center of the guanidino group of Arg. In the case of R346X, any substitution except lysine would result in a side chain that is significantly shorter and non-cationic, thus dissolving the interactions between N450 or other common substitutions at that position.

Other combinations have been exceedingly rare so far, and seen only in cryptic lineages (e.g., F486P and K444 mutations), but no steric justifications can be found for them.

### Epistasis

While the focus so far has been mostly on the Spike protein, it is likely that convergent evolution is acting on genes other than Spike. Given that the Spike protein is the best protective antigen for both infection and vaccines, mutations in other genes are more likely to provide fitness advantages if they affect Spike expression. E.g., ORF8 limits the amount of Spike proteins that reaches the cell surface and is incorporated into virions, reducing recognition by anti-SARS-CoV-2 antibodies[30]. ORF8 has accordingly been target of convergent evolution in Omicron (e.g., ORF8:S667F in BR.2.1, ORF8:G8x in XBB.1) and in SARS-related coronaviruses[31].

Other genes whose roles in Spike modulation are not clear are also converging, such as ORF1b:T1050, found in many BA.5.2.* sublineages, and XBE (T1050N) as well as XBC.* (T1050I).

### Selective pressures from therapeutics targeting the Spike protein

There is a theoretical concern that, in addition to vaccines- and infection-elicited immunity, selective pressure by prophylactic and therapeutic anti-Spike monoclonal antibodies (mAb), can contribute to the emergence of novel SARS-CoV-2 sublineages [32]. While selective pressures are likely to generate many different mutants, a very few of those emerging sublineages could be fit enough to compete with the lineages that are dominating at that time to become locally or globally dominant.

While spontaneous evolution can occur in the absence of selective pressures due to the intrinsic genomic MR (see section above), extended half-life mAbs (such as Evusheld™) administered for pre-exposure prophylaxis or therapy to chronically infected immunocompromised patients at subneutralizing concentrations provide ideal conditions to facilitate the emergence of mutants[33], for these patients often cannot clear the infection and have high viral loads. Establishing a cause-effect relationship is difficult, but intra-host evolution studies provide a highly suggestive temporal association[34]. mAbs have come of age since the advent of the SARS-CoV-2 Delta VOC, but because of the resistance of Omicron to most authorized mAbs, their use since Spring 2022 has been largely limited to Evusheld™ (for which cilgavimab was the only ingredient with residual activity) and bebtelovimab.

We know from *in vitro* deep mutational scanning studies the exact mutations that cause resistance to each mAb. S:F486X mutations impart resistance to tixagevimab, S:R346X, S:K444X and S:S494X mutations impart resistance to cilgavimab, while S:K444X mutations impart resistance to bebtelovimab (Table 1). We recently noted an increase in the circulation of Omicron sublineages associated with S:R346X mutations, and wondered whether this could partly be the result of selective pressure with Evusheld™. We compared the prevalence of R346X mutations in countries with high versus low usage of Evusheld™ (France vs. UK) or bebtelovimab (USA vs. UK) (Figure 8). UK also represents an ideal control because of its very high SARS-CoV-2 genome sequencing rate. We discuss these 2 scenarios in details below.

**Table 1.**
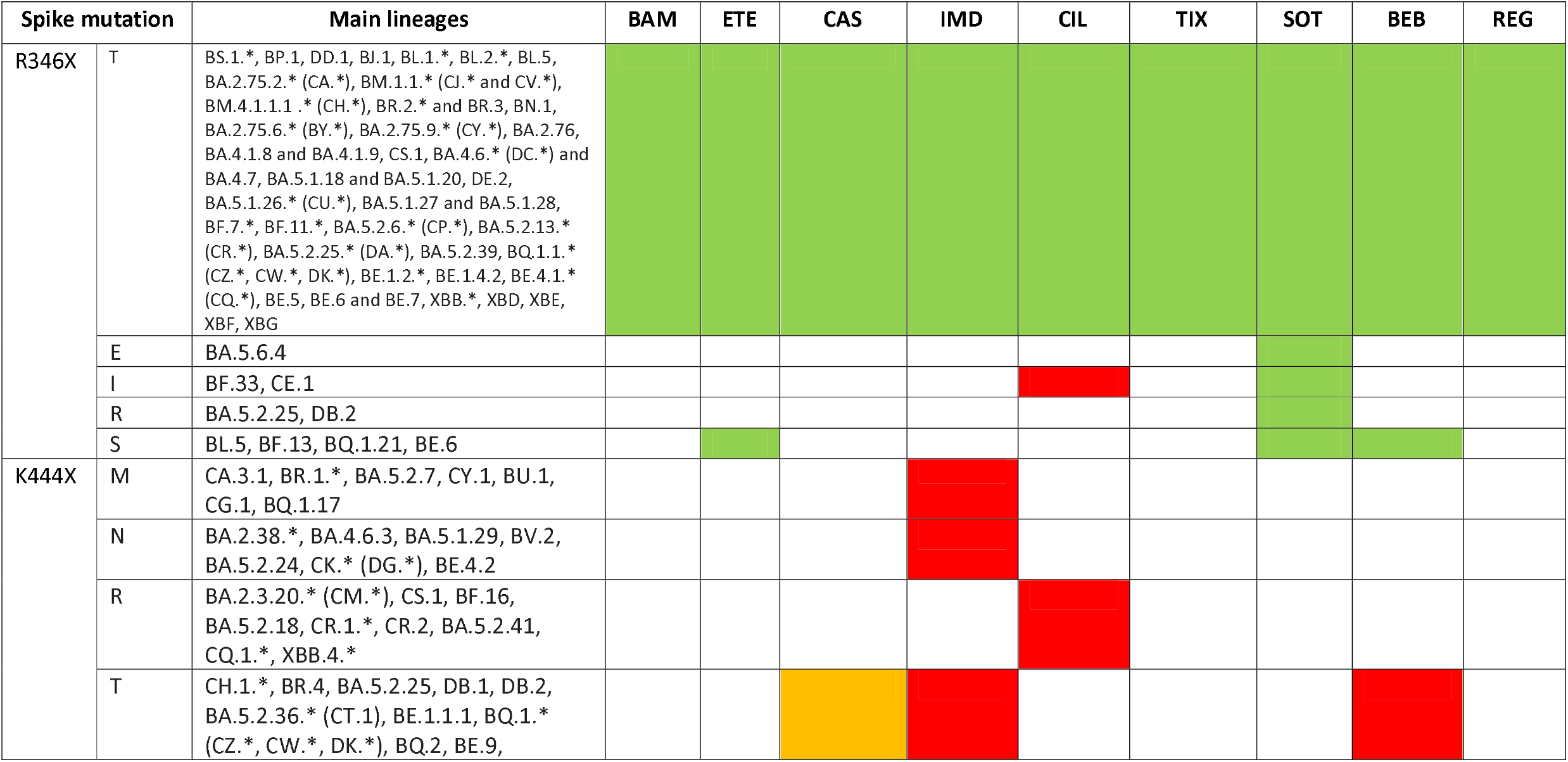
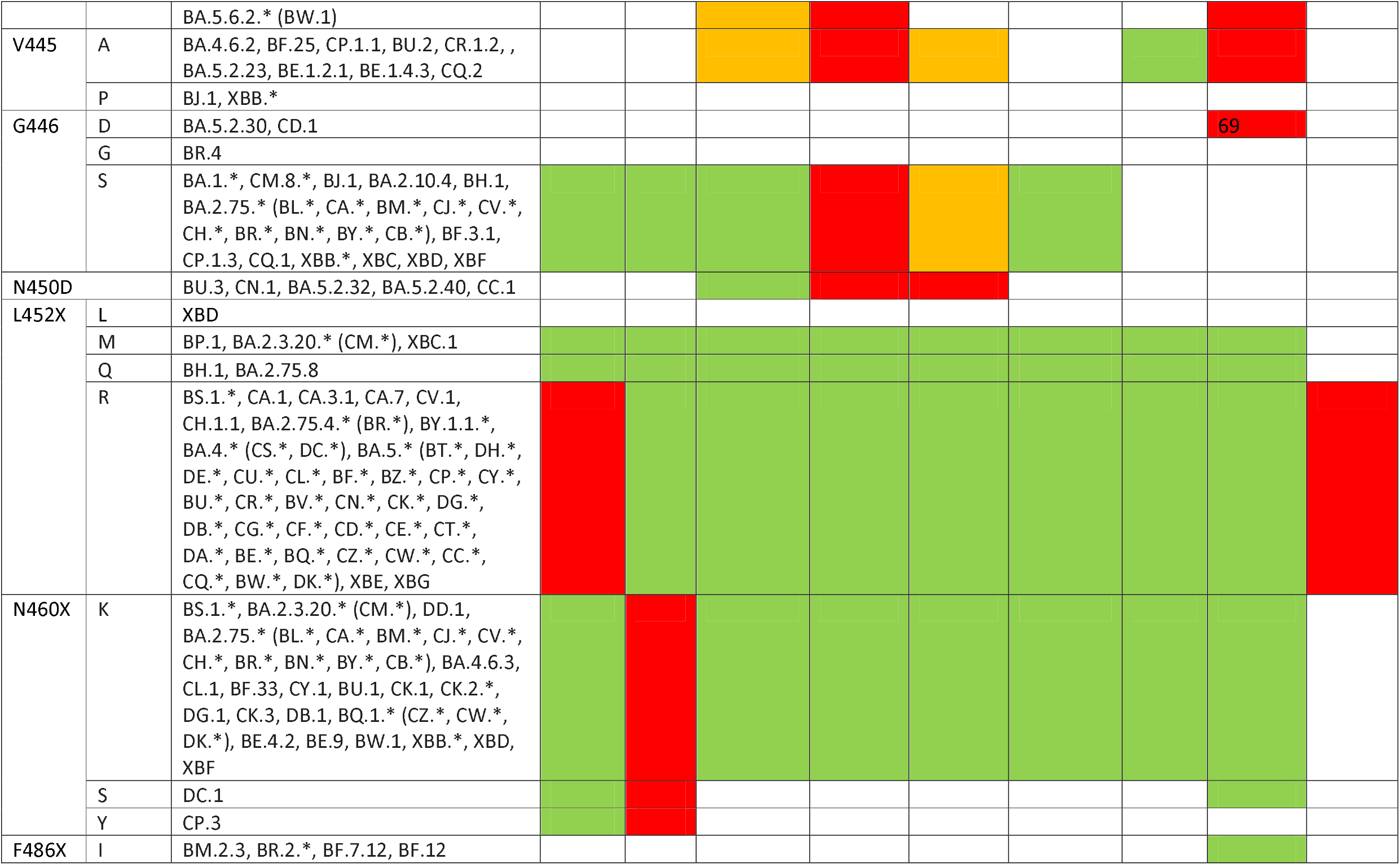
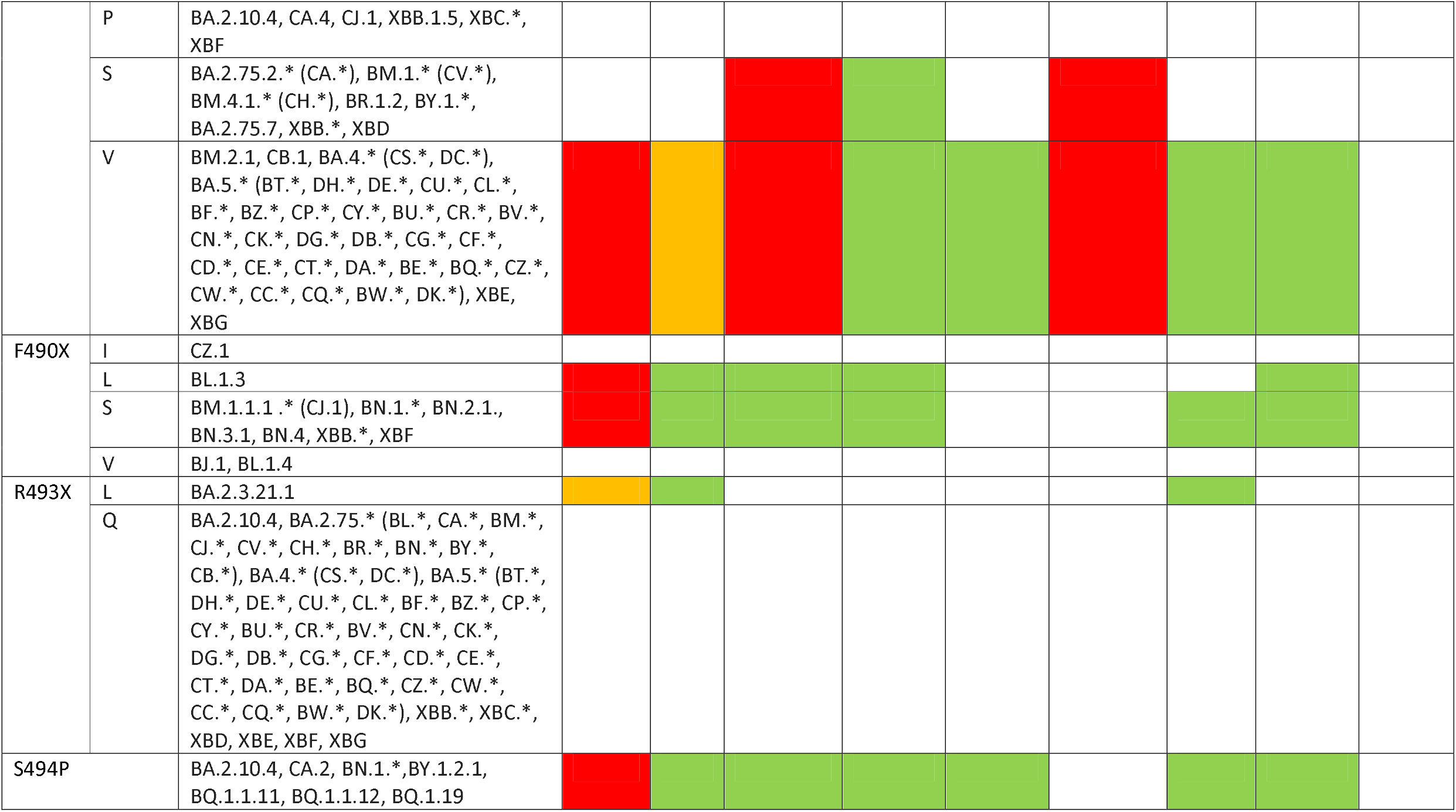
Heatmap of selected Spike RBD mutations in Omicron sublineages and their impact on authorized therapeutic anti-Spike mAbs. BAM: bamlanivimab; ETE: etesevimab; CAS: casirivimab; IMD: imdevimab; TIX: tixagevimab; CIL: cilgavimab; SOT: sotrovimab; BEB: bebtelovimab; REG: regdanvimab. Data sourced from the Stanford University Coronavirus and Antiviral Resistance Database (accessed online at https://covdb.stanford.edu/search-drdb on November 30, 2022). Green means fold-reductions < 5; Orange means fold-reduction 5-100 ; Red means fold-reduction in IC_50_ > 100 compared to wild-type; blank means no data available.

**Figure 8.**
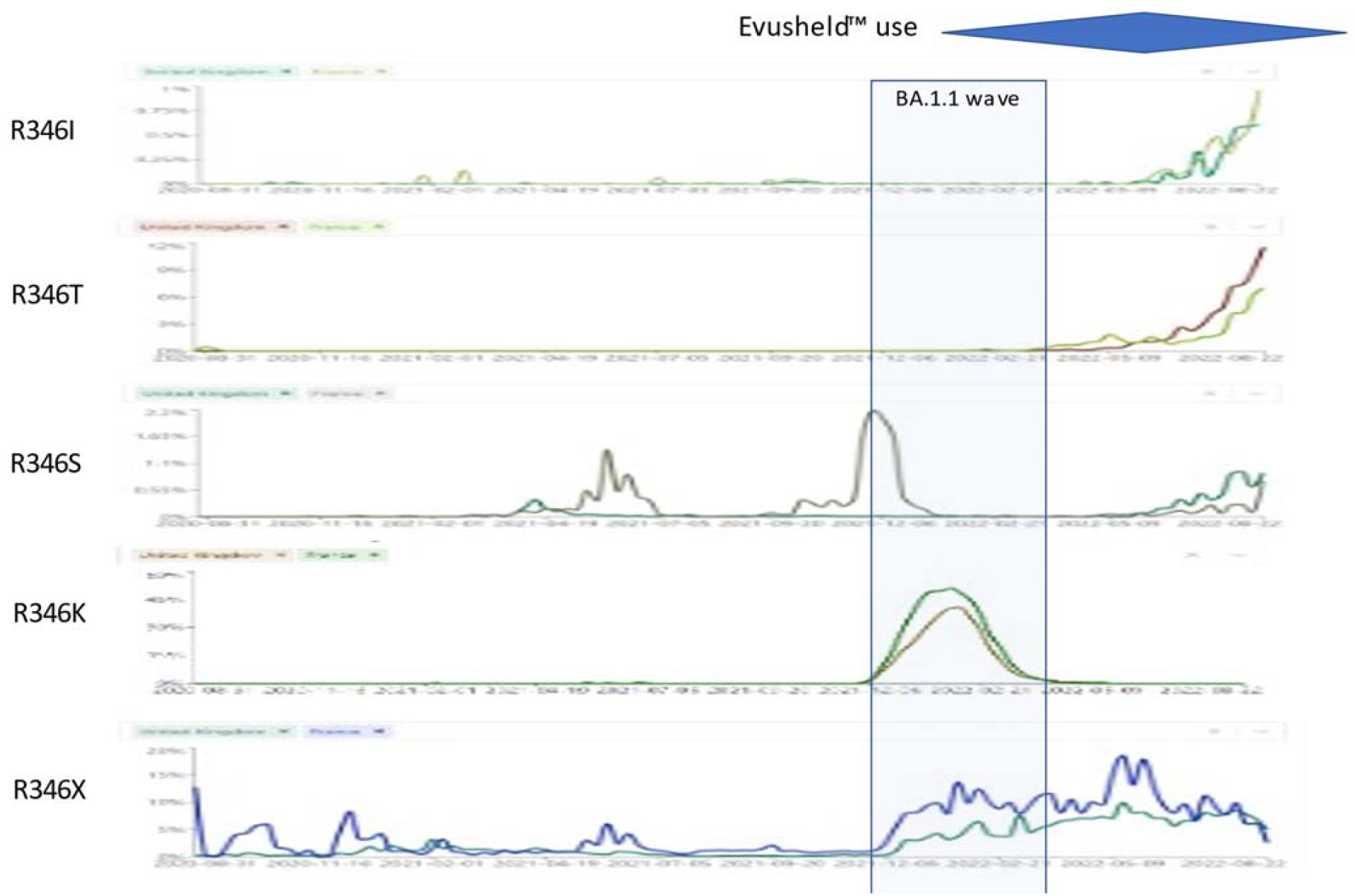
Prevalence of S:R346X mutations in the period 2020-08-27 – 2022-09-13 in UK versus France. Sourced from https://cov-spectrum.org on November 26, 2022. The blue area represents trends in Evusheld™ prescriptions in France and many other countries (but not UK).

### S:R346X

Different mutations can affect the R346 residue. R346G has been selected *in vitro* by cilgavimab+tixagevimab[35]. R346S occurred *in vitro* after 12 weeks of propagating SARS-CoV-2 in the presence of sotrovimab, and before the other epitope mutation (P337L) which leads to sotrovimab resistance [36]. R346I has been selected *in vitro* under the selective pressure from cilgavimab [37,38]. Lee *et al* reported mutually exclusive substitutions at residues R346 (R346S and R346I) and E484 (E484K and E484A) of Spike protein and continuous turnover of these substitutions in 2 immunosuppressed patients[39]. Unfortunately, *in vivo* selection evidences are so far available for sotrovimab[40] but not for Evusheld™. It should be anyway noted that R346T[41,42] and R346I[43] have been reported to spontaneously develop and fix in 3 IC patients without any selective pressure.

While R346K was associated with the BA.1.1 wave (see Figure 8), the plethora of different Omicron sublineages that showed convergent evolution towards R346I, R346S or R346T is of concern.

- R346K (previously seen only in VOC Mu/B.1.621 [44]) occurred exclusively in BA.1.1, a sublineage that disappeared since May 2022, where it affected the interaction network in the BA.1.1 RBD/hACE2 interface through long-range alterations and contributes to the higher hACE2 affinity of the BA.1.1 RBD than the BA.1 RBD [45], and had increased resistance against Evusheld™[46] and sotrovimab[47]. Only STI-9167 remained effective among the mAbs[48]. Beta+R346K, which was identified in the Philippines in August 2021, exhibited the highest resistance to 2 BNT61b2 doses-elicited sera among the tested VOCs[49]. After BA.1.1, R346K has not been detected worldwide in any sublineage.
- R346I occurs in more than 40 different Omicron sublineages, but it is most represented in BA.5.9 (38%), BA.4.1 (5%), BA.5.9 (4%), but also occurred in AY.39 (14%);
- R346S (previously seen only in a C.36.3 sublineage from Italy[50] (30.8%), occurs in more than 40 different Omicron sublineages but it is most represented in B.1.640.1 (18%), and in a few Delta sublineages (<2%)) occurs nowadays in BA.4.7 (13%), BA.5.2.1 (8.22%), BA.4 (2.8%).
- R346T occurs in more than 96 different Omicron sublineages, but it is mostly represented in BA.4.6 (44%), BA.5.2.1 (13%), BA.2 (8%), BA.2.74 (3%), BA.2.76 (12%), BA.4.1 (2.3%). In addition, it is a hallmark mutation of BA.1.23, BA.2.9.4, BL.1, BA.2.75.2, BA.2.80, BA.2.82, BA.4.1.8, BF.7 and BF.11. BA.4.6, BA.4.7, and BA.5.9 displayed higher humoral immunity evasion capability than BA.4/BA.5, causing 1.5 to 1.9-fold decrease in NT_50_ of the plasma from BA.1 and BA.2 breakthrough-infection convalescents compared to BA.4/BA.5. Importantly, plasma from BA.5 breakthrough-infection convalescents also exhibits significant neutralization activity decrease against BA.4.6, BA.4.7, and BA.5.9 than BA.4/BA.5, showing on average 2.4 to 2.6-fold decrease in NT_50_. R346S causes resistance to class 3 antibodies: bebtelovimab remains potent, while Evusheld™ is completely escaped by these subvariants [51].

### S:K444X

The K444E/R mutations were reported *in vitro* after selection with cilgavimab[38]. Resistance studies with bebtelovimab selected the K444T escape mutations for BA.2[52]. Ortega *et al* found that K444R (previously found in the Beta VOC[53]), K444Q, and K444N mutations can change the virus binding affinity to the ACE2 receptor[54]. Weisblum *et al* found that K444R/Q/N occurs after exposure to convalescent plasma[55]. Among largely diversified VOCs such as Delta, S:K444N was associated with reduced remdesivir binding and increased mortality[56].

## Conclusions

The convergent evolution of Omicron sublineages appears to reflect the selective pressure exerted by previous infection- or vaccine-elicited immunity. Vaccines and perhaps antibody therapeutics have without doubt saved an untold number of lives but also likely altered the natural evolutionary trajectory of the virus. While other viruses such as influenza and HIV routinely produced new variants because of their mutagenicity, the scale at which SARS-CoV-2 has spun off new variants and lineages appears unprecedented in modern virology history. The SARS-CoV-2 vaccines reduce severe disease and mortality but do not confer sufficient immunity to prevent re-infection with viral replication in vaccinated hosts. Hence, we have the unusual situation of viral replication in immune hosts where the immune system is placing evolutionary pressure on the virus to select variants that can escape vaccine-elicited immunity in addition to infection-elicited immunity. Whether this rapid evolutionary trajectory is the result of viral replication properties, replication in immune hosts or both is unknown but conditions present in the past year of the pandemic have produced a remarkable natural experiment in viral evolution for which we cannot discern its conclusion.

Insights from structural biology has shown how some mutations are mutually exclusive, which could help the design of next-generation vaccines. But the latter could reset the run for immune escape, perpetuating the never-ending game of host and pathogen. Viral recombination[57] (more than 50 lineages censed at the time of writing, with both simple and complex variants[58]) and sudden reemergence of former VOCs[59] have to be considered as further drivers for evolutionary saltation.

In this setting, polyclonal passive immunotherapies (such as plasma from convalescent and vaccinated donors[60,61]) appear more escape-resistant than monoclonal antibodies[62-65], and combo therapies should be urgently investigated and deployed in vulnerable populations, such as IC patients[66].

## Abbreviations

IC: immunocompromised
MR: mutation rate
RBM: receptor-binding motif
VOC: variant of concern.

## Acknowledgments

we are grateful to Cornelius Roemer (Biozentrum - Universität Basel, Switzerland), Thomas P Peacock (Imperial College, UK), independent researchers Ryan Hisner and Federico Gueli, and the Twitter user @FanDorop for discussions about convergent evolution.

---

**Supplementary Table 1.**
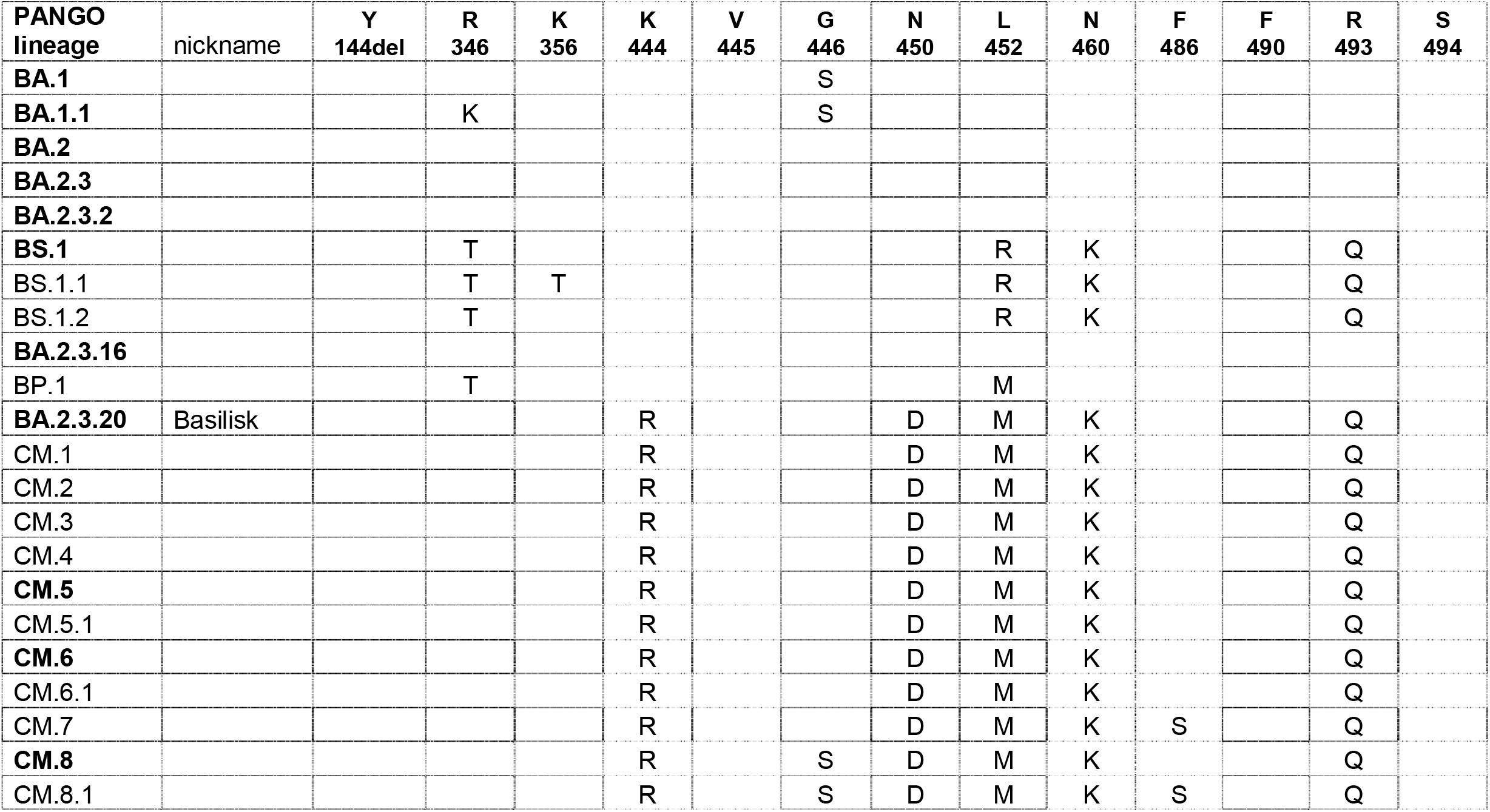
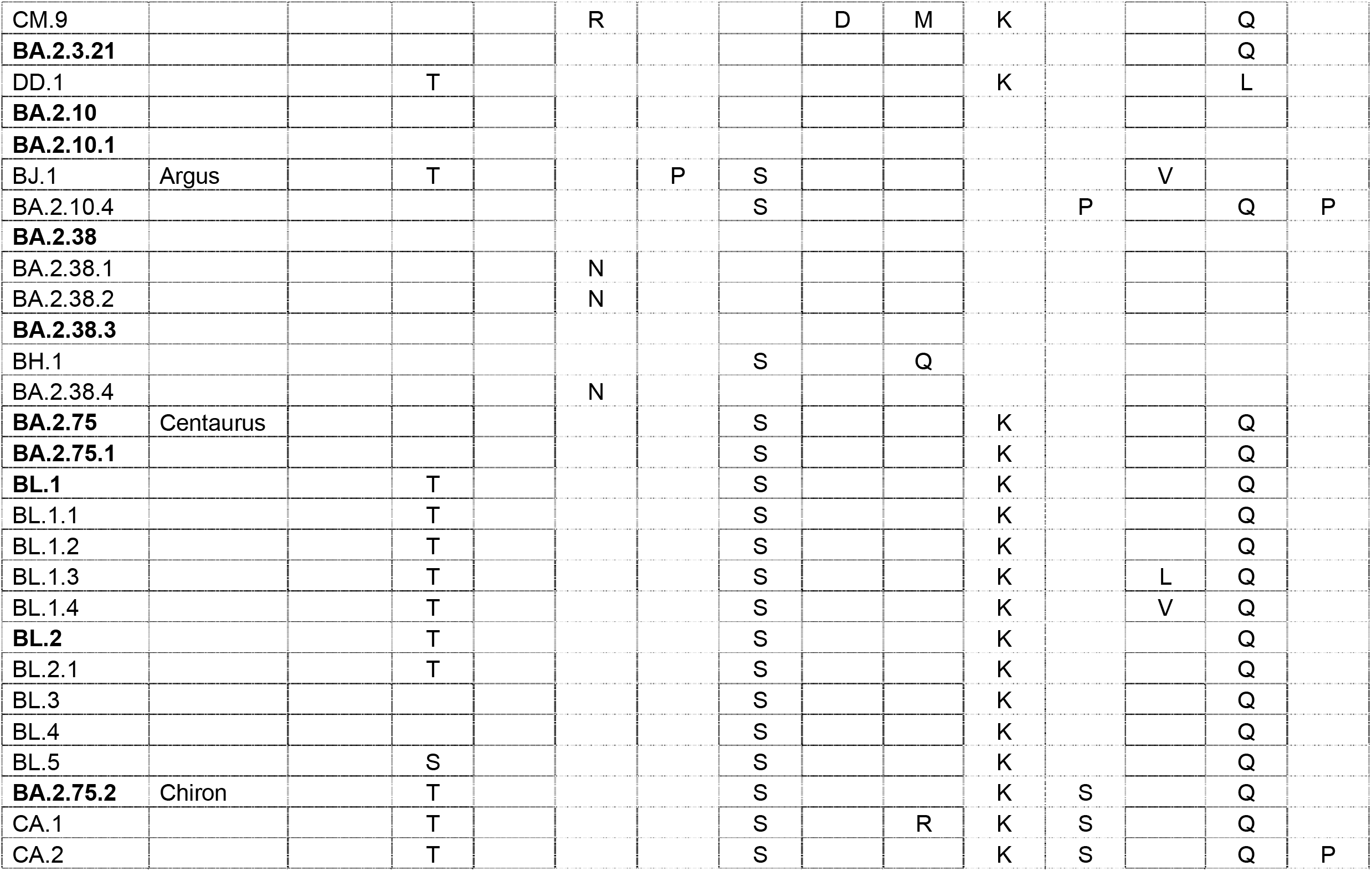
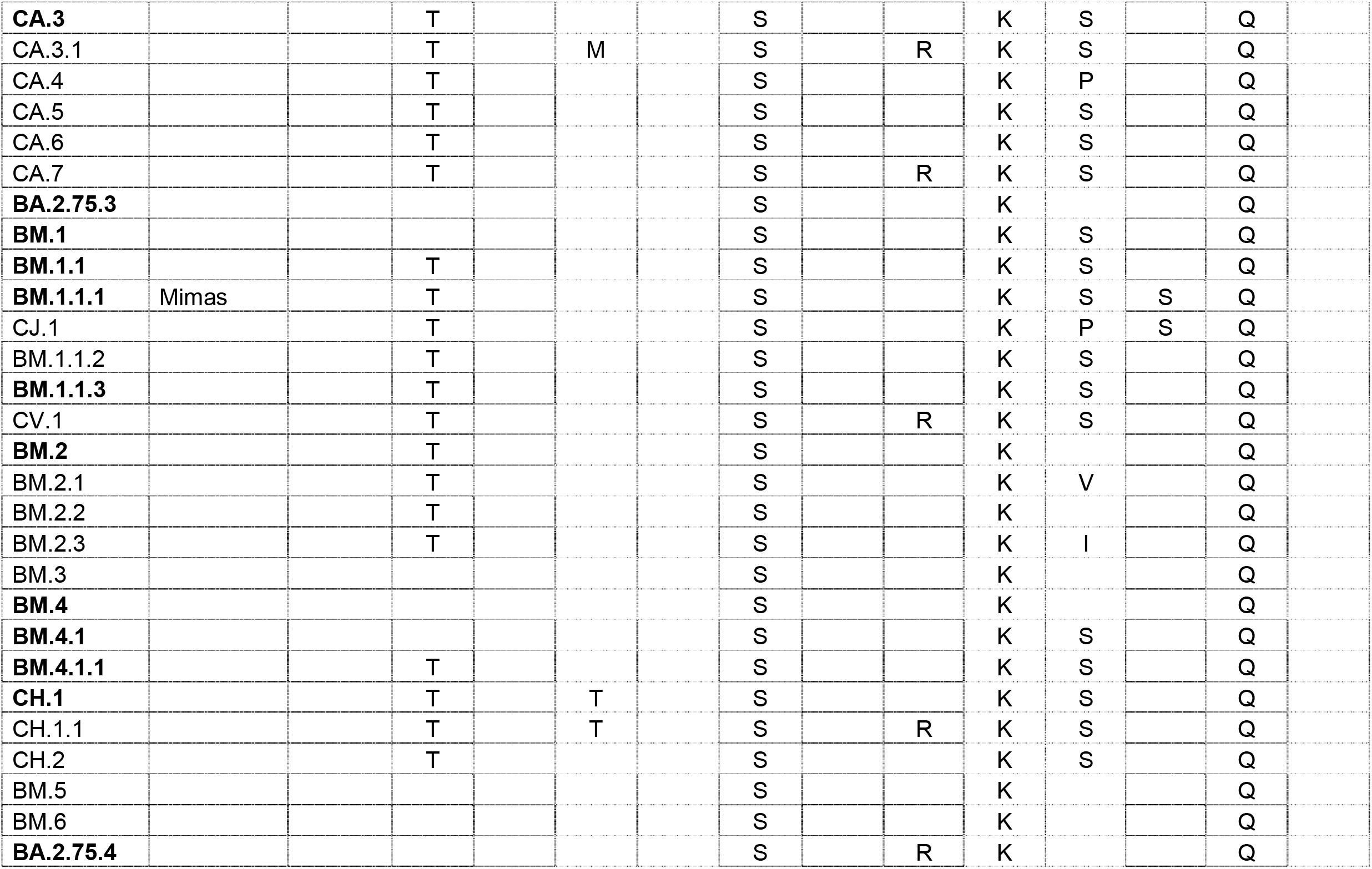
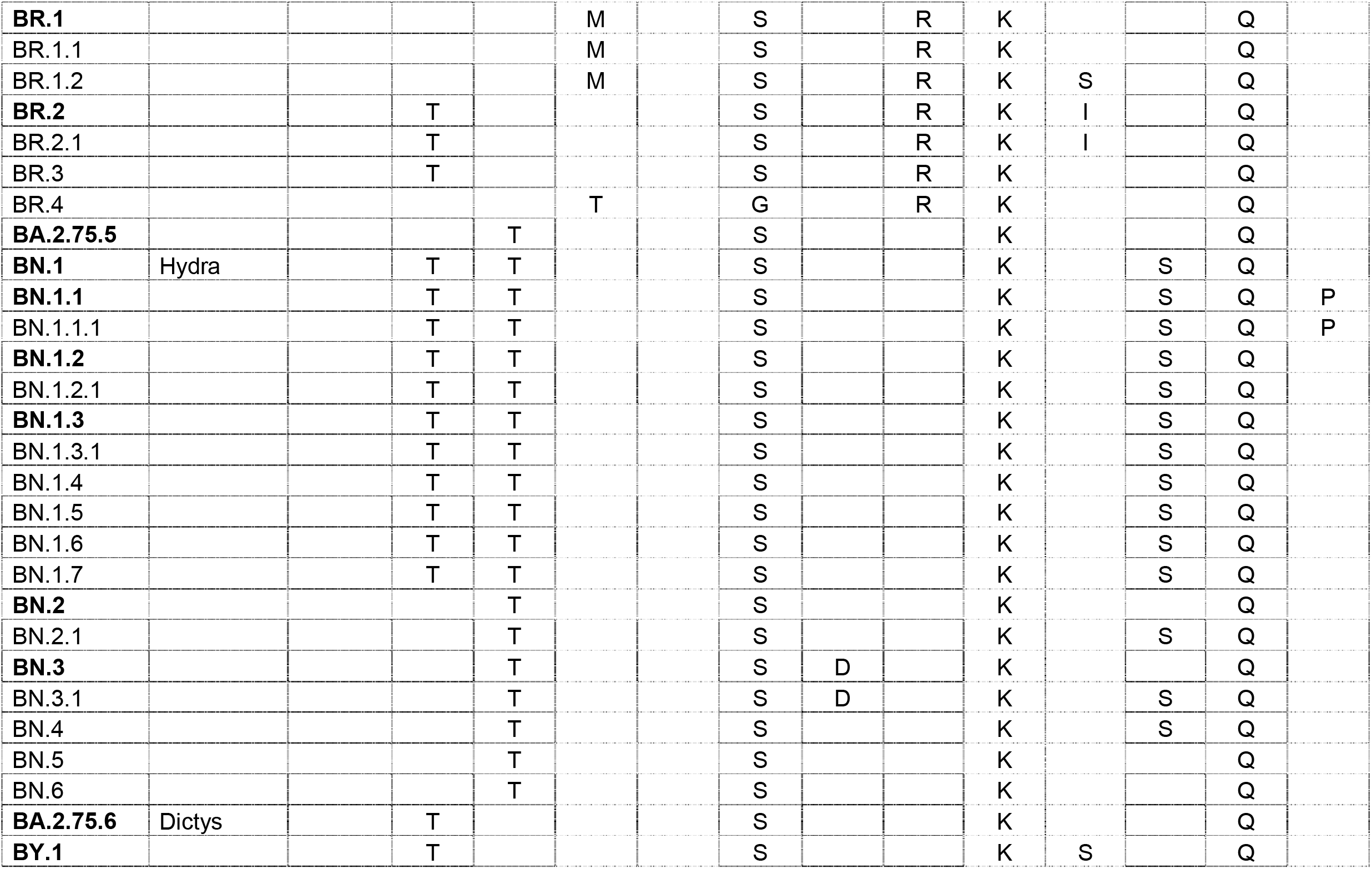
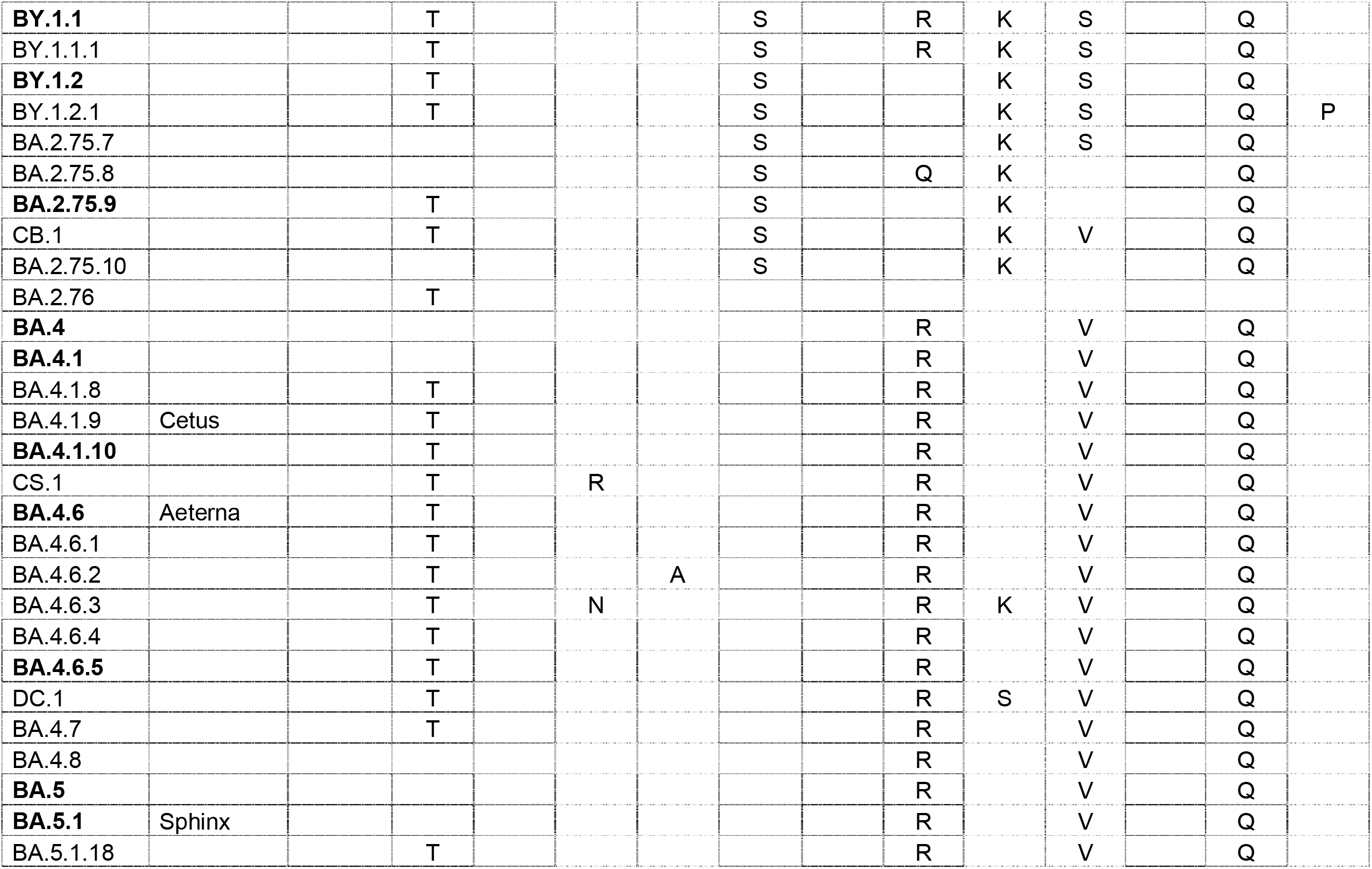
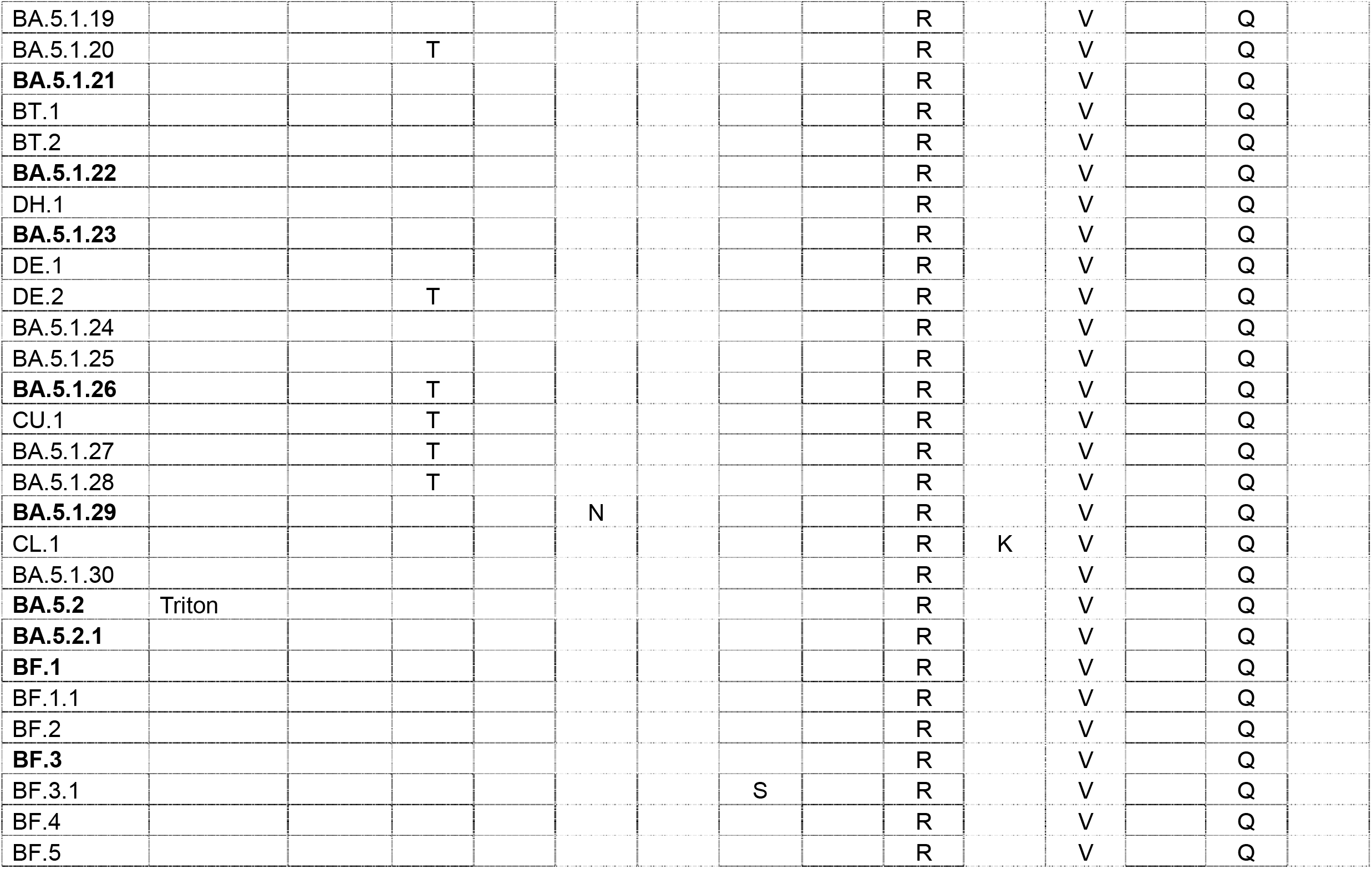
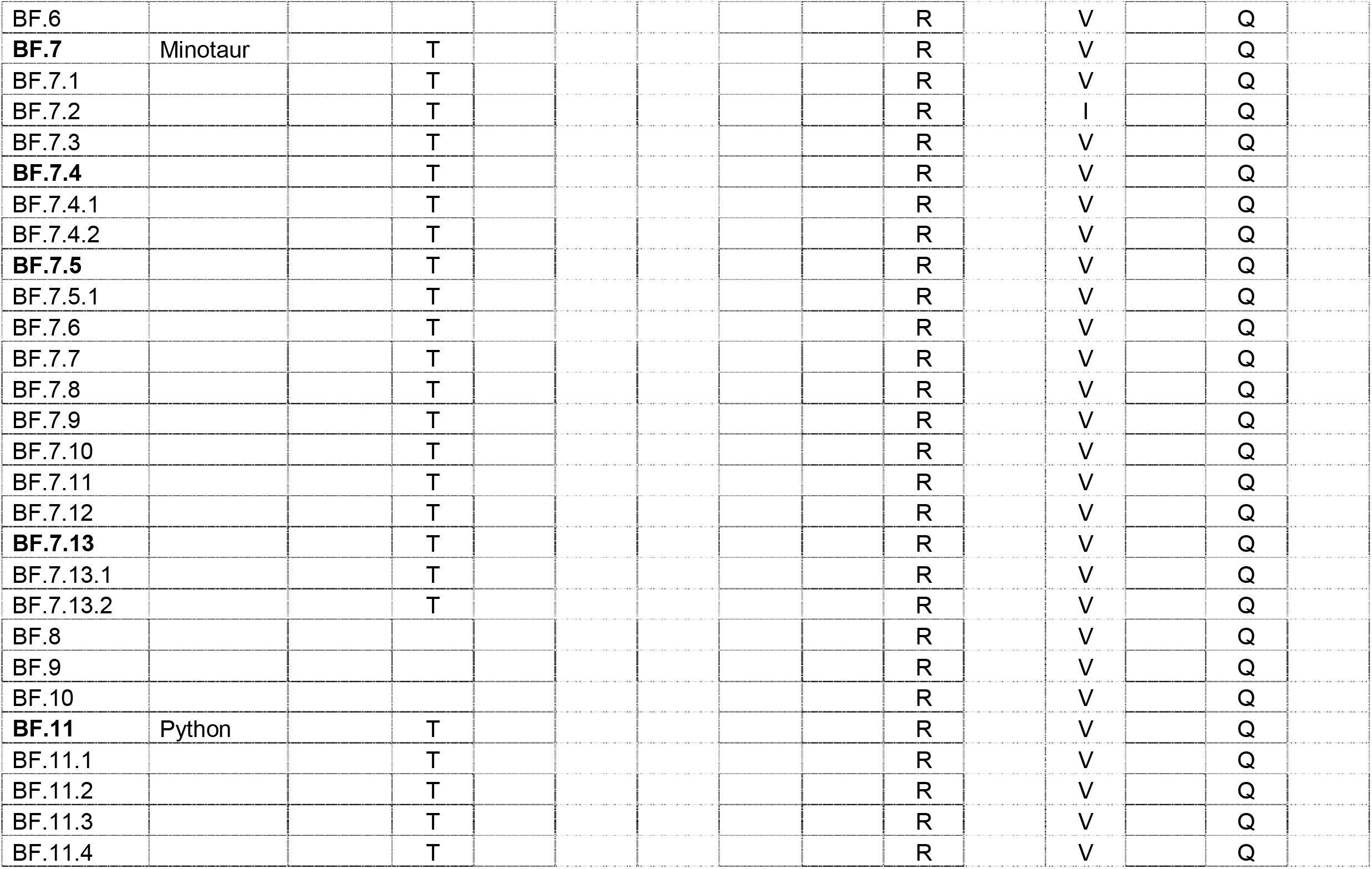
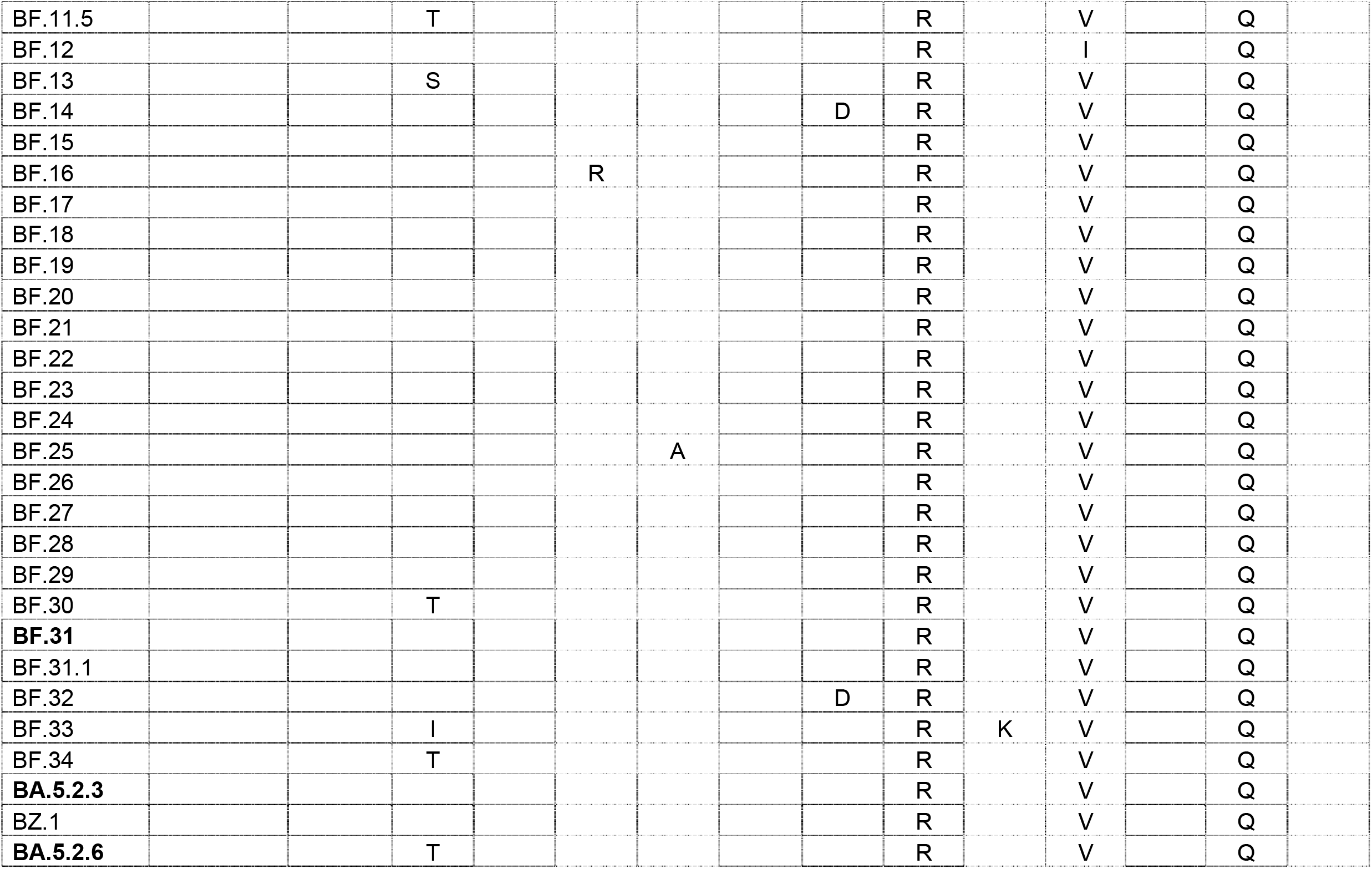
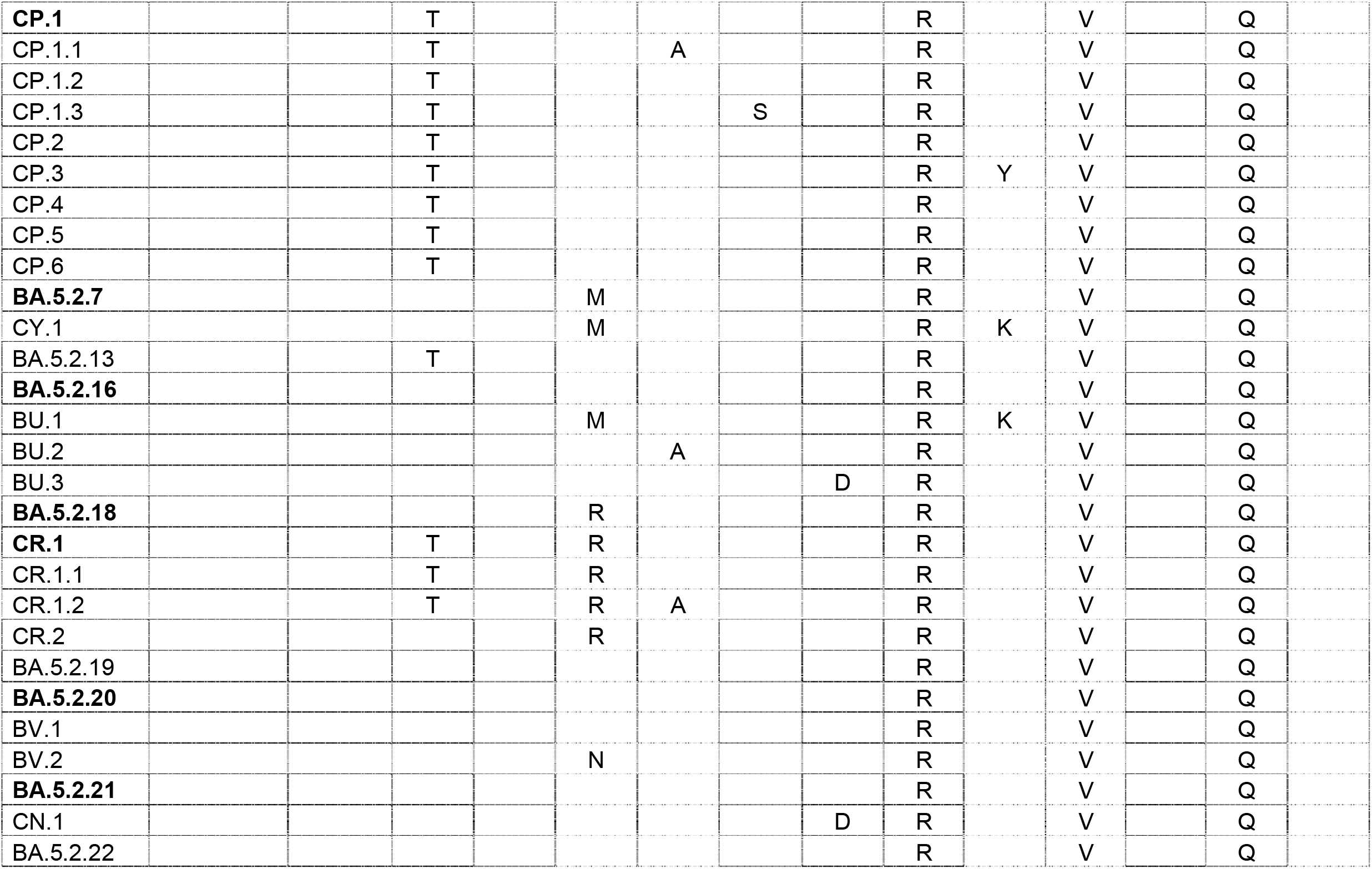
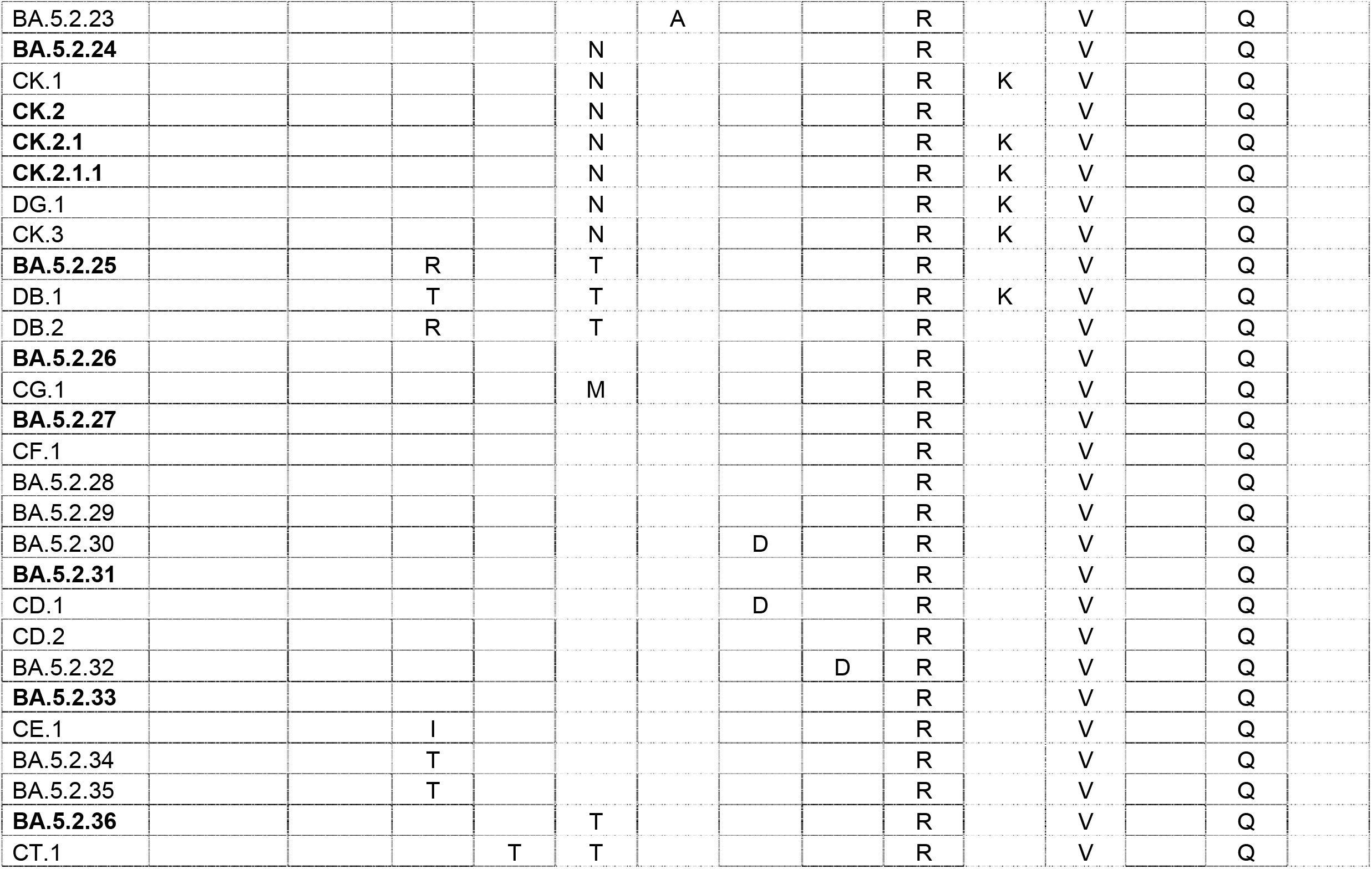
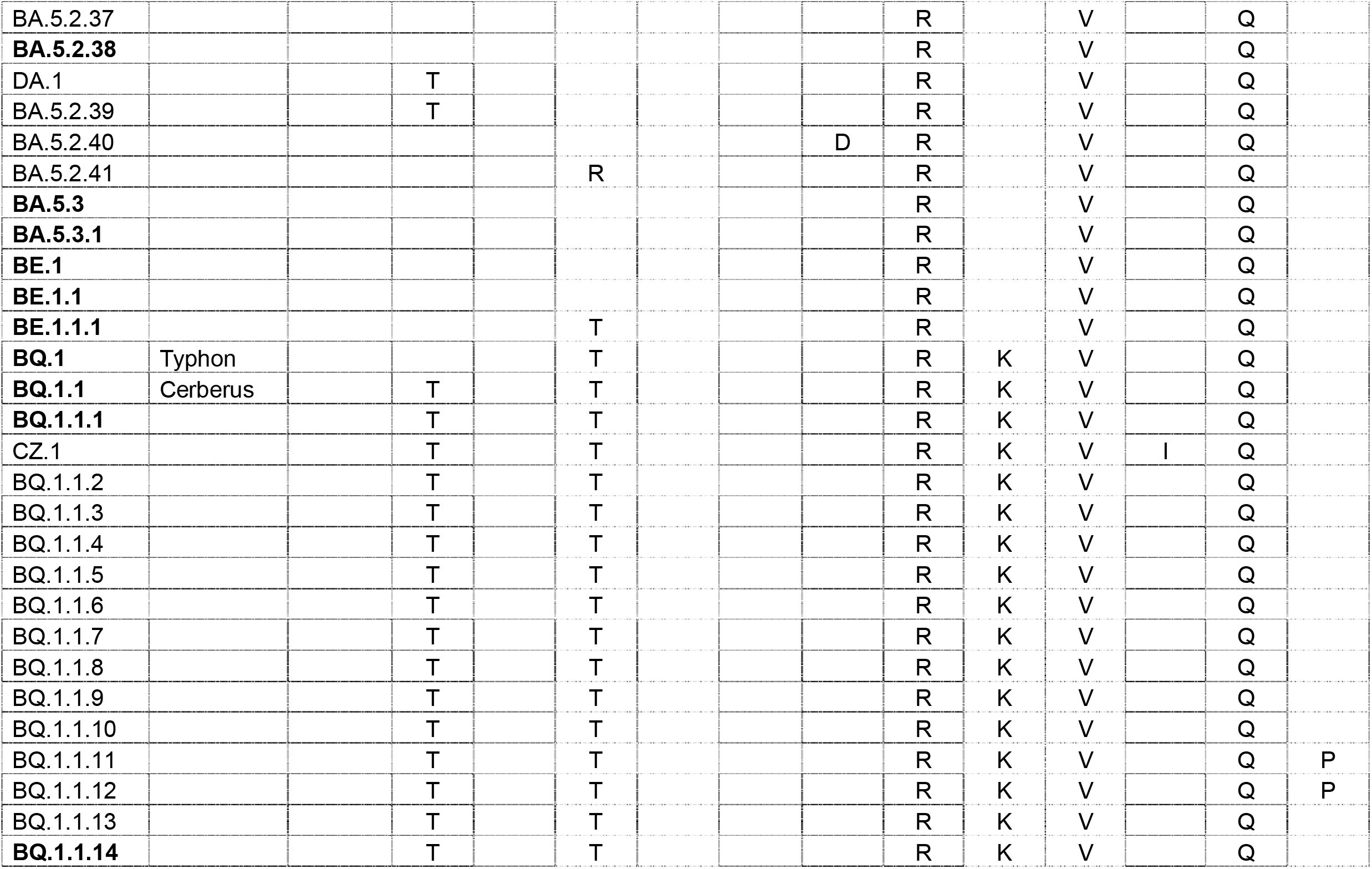
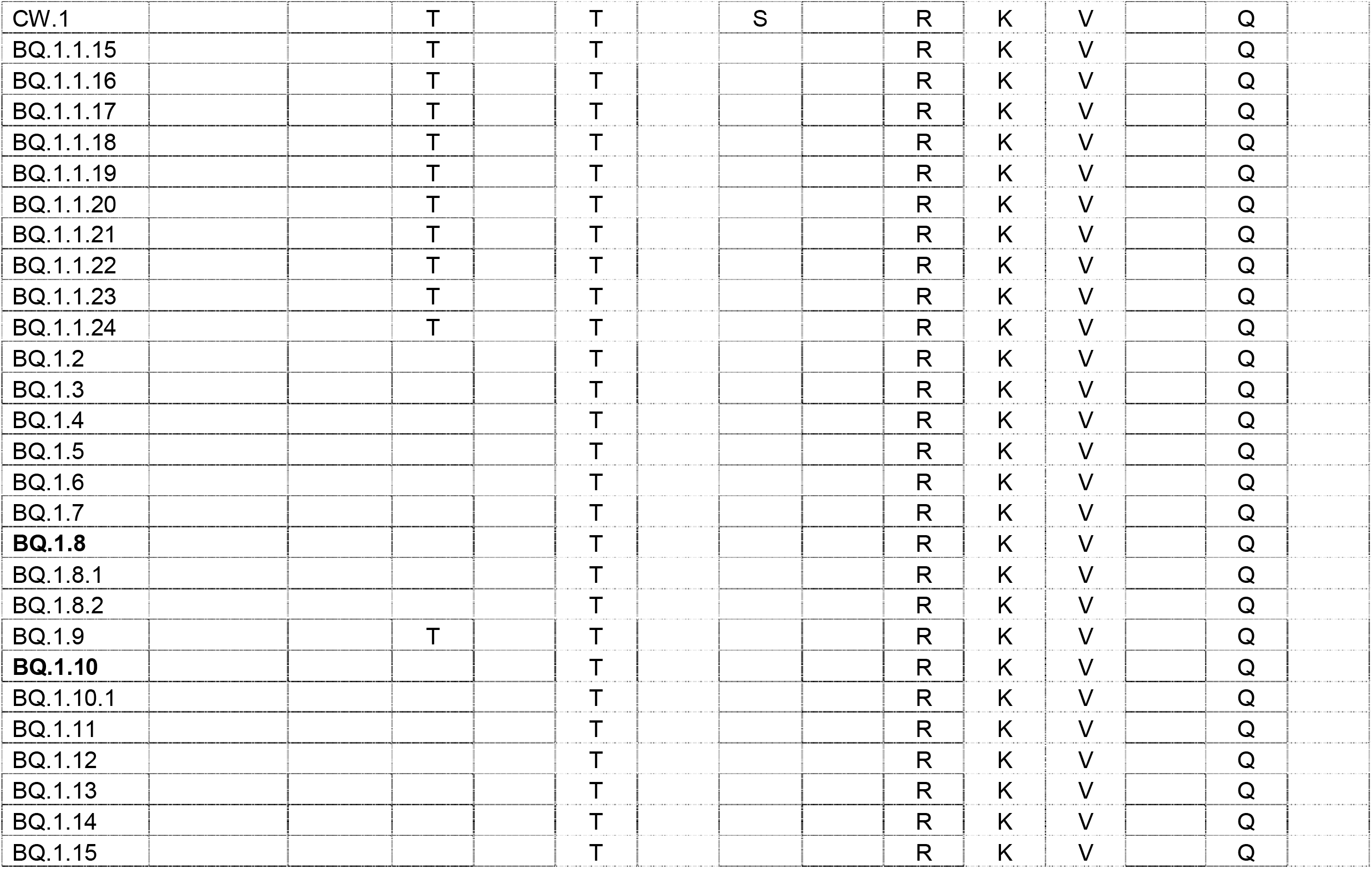
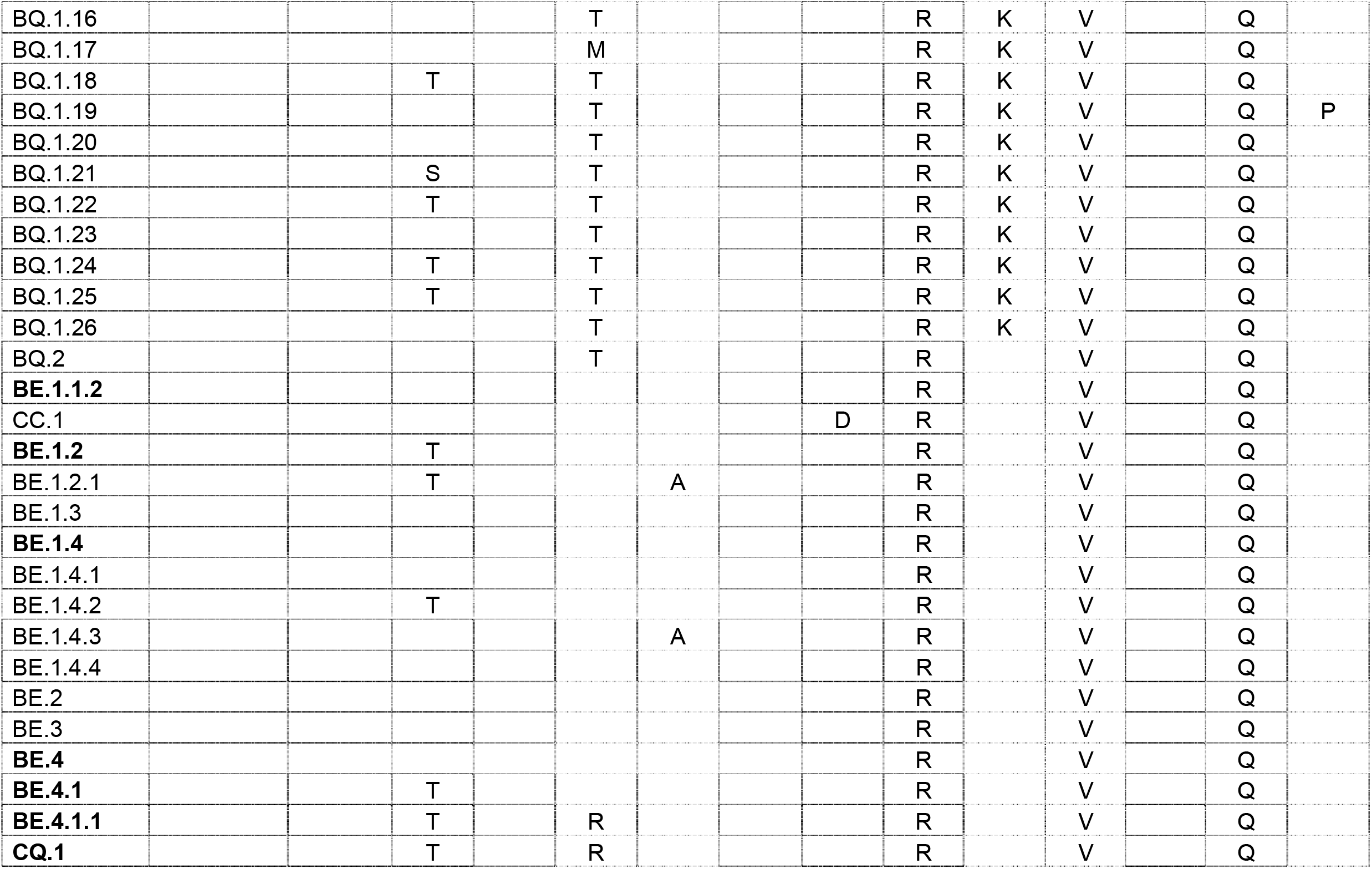
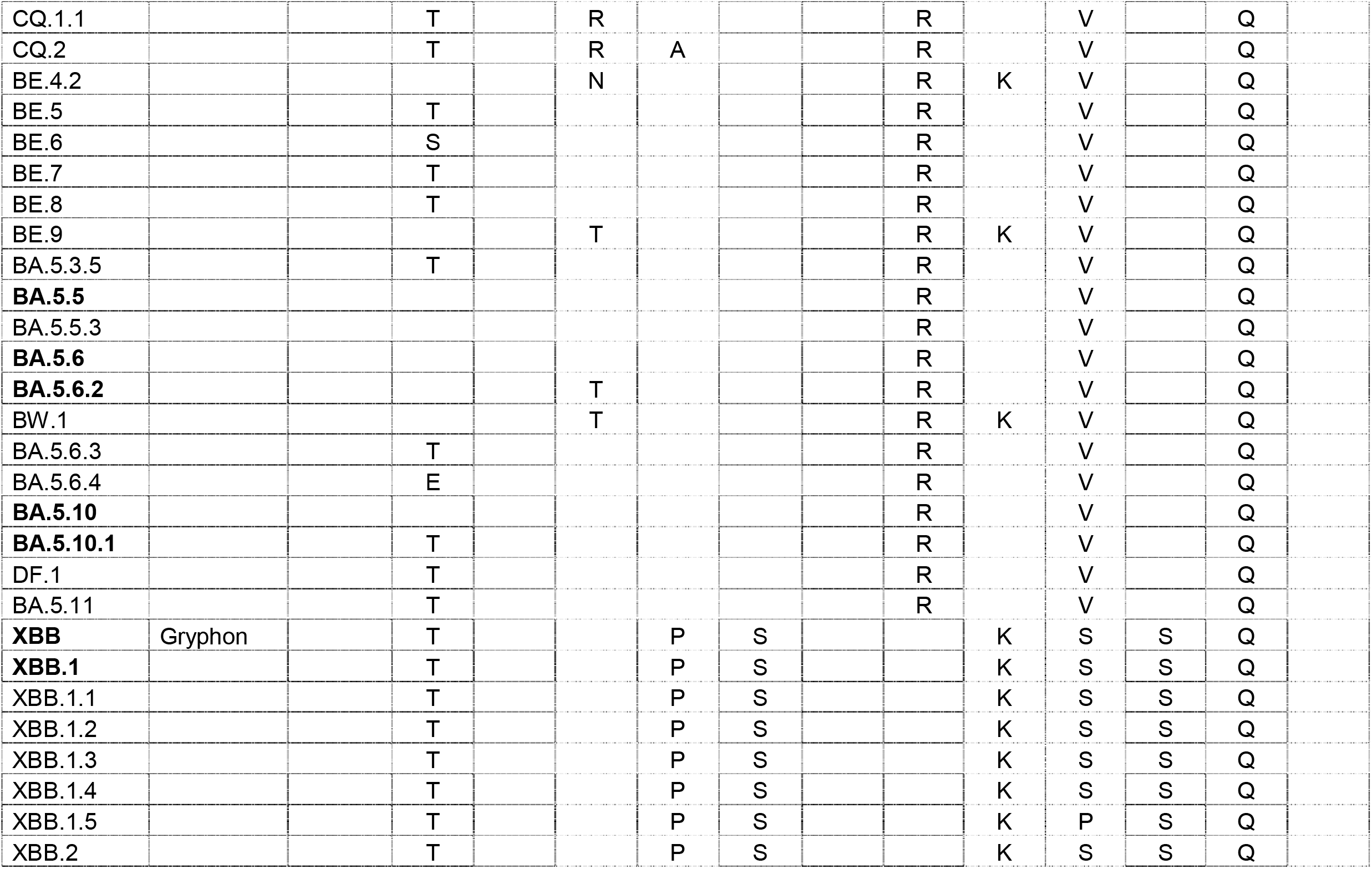
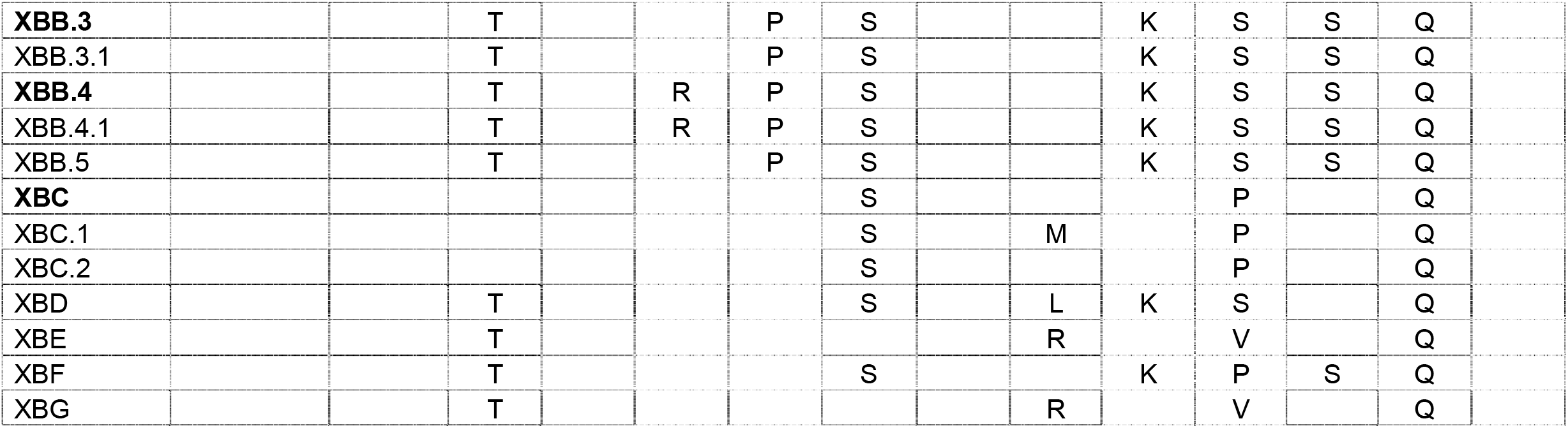
Occurrence of selected amino acid mutations associated with immune escape within the Spike protein of SARS-CoV-2 in BA.2 and BA.4/5 Omicron sublineages. Modified from https://docs.google.com/spreadsheets/d/1OTWogpyvWNTlK0ww7TlDcI4l_SkZt3Z8nYiSp2YARys/edit?usp=sharing

## References

1. Daily new confirmed COVID-19 deaths per million people. Our world in data. Accessed online at https://ourworldindata.org/explorers/coronavirus-data-explorer?zoomToSelection=true&time=2020-03-01..latest&facet=none&pickerSort=asc&pickerMetric=location&hideControls=false&Metric=Confirmed+deaths&Interval=7-day+rolling+average&Relative+to+Population=true&Color+by+test+positivity=false&country=~OWID_WRL on December 1, 2022.

2. Viana, R.; Moyo, S.; Amoako, D.G.; Tegally, H.; Scheepers, C.; Althaus, C.L.; Anyaneji, U.J.; Bester, P.A.; Boni, M.F.; Chand, M.; et al. Rapid epidemic expansion of the SARS-CoV-2 Omicron variant in southern Africa. Nature 2022, 603, 679–686, doi:10.1038/s41586-022-04411-y.

3. Harari, S.; Tahor, M.; Rutsinsky, N.; Meijer, S.; Miller, D.; Henig, O.; Halutz, O.; Levytskyi, K.; Ben-Ami, R.; Adler, A.; et al. Drivers of adaptive evolution during chronic SARS-CoV-2 infections. Nat Med 2022, 28, 1501–1508, doi:10.1038/s41591-022-01882-4.

4. Kemp, S.A.; Collier, D.A.; Datir, R.P.; Ferreira, I.A.T.M.; Gayed, S.; Jahun, A.; Hosmillo, M.; Rees-Spear, C.; Mlcochova, P.; Lumb, I.U.; et al. SARS-CoV-2 evolution during treatment of chronic infection. Nature 2021, 592, 277–282, doi:10.1038/s41586-021-03291-y.

5. Wilkinson, S.A.J.; Richter, A.; Casey, A.; Osman, H.; Mirza, J.D.; Stockton, J.; Quick, J.; Ratcliffe, L.; Sparks, N.; Cumley, N.; et al. Recurrent SARS-CoV-2 mutations in immunodeficient patients. Virus Evol 2022, 8, veac050, doi:10.1093/ve/veac050.

6. Focosi, D. Molnupiravir: From Hope to Epic Fail? 2022, 14, 2560.

7. Smith, D.J.; Lapedes, A.S.; de Jong, J.C.; Bestebroer, T.M.; Rimmelzwaan, G.F.; Osterhaus, A.D.; Fouchier, R.A. Mapping the antigenic and genetic evolution of influenza virus. Science 2004, 305, 371–376, doi:10.1126/science.1097211.

8. Eguia, R.T.; Crawford, K.H.D.; Stevens-Ayers, T.; Kelnhofer-Millevolte, L.; Greninger, A.L.; Englund, J.A.; Boeckh, M.J.; Bloom, J.D. A human coronavirus evolves antigenically to escape antibody immunity. PLoS Pathog 2021, 17, e1009453, doi:10.1371/journal.ppat.1009453.

9. Niesen, M.; Anand, P.; Silvert, E.; Suratekar, R.; Pawlowski, C.; Ghosh, P.; Lenehan, P.; Hughes, T.; Zemmour, D.; OHoro, J.C.; et al. COVID-19 vaccines dampen genomic diversity of SARS-CoV-2: Unvaccinated patients exhibit more antigenic mutational variance. 2021, 2021.2007.2001.21259833, doi:10.1101/2021.07.01.21259833 %J medRxiv.

10. Yeh, T.-Y.; Contreras, G.P. Full vaccination is imperative to suppress SARS-CoV-2 delta variant mutation frequency. 2021, 2021.2008.2008.21261768, doi:10.1101/2021.08.08.21261768 %J medRxiv.

11. Ruis, C.; Peacock, T.P.; Polo, L.M.; Masone, D.; Alvarez, M.S.; Hinrichs, A.S.; Turakhia, Y.; Cheng, Y.; McBroome, J.; Corbett-Detig, R.; et al. Mutational spectra distinguish SARS-CoV-2 replication niches. 2022, 2022.2009.2027.509649, doi:10.1101/2022.09.27.509649 %J bioRxiv.

12. Bloom, J.D.; Beichman, A.C.; Neher, R.A.; Harris, K. Evolution of the SARS-CoV-2 mutational spectrum. 2022, 2022.2011.2019.517207, doi:10.1101/2022.11.19.517207 %J bioRxiv.

13. V’Kovski, P.; Kratzel, A.; Steiner, S.; Stalder, H.; Thiel, V. Coronavirus biology and replication: implications for SARS-CoV-2. Nature reviews. Microbiology 2021, 19, 155–170, doi:10.1038/s41579-020-00468-6.

14. Sadler, H.A.; Stenglein, M.D.; Harris, R.S.; Mansky, L.M. APOBEC3G contributes to HIV-1 variation through sublethal mutagenesis. Journal of virology 2010, 84, 7396–7404, doi:10.1128/jvi.00056-10.

15. De Maio, N.; Walker, C.R.; Turakhia, Y.; Lanfear, R.; Corbett-Detig, R.; Goldman, N. Mutation Rates and Selection on Synonymous Mutations in SARS-CoV-2. Genome biology and evolution 2021, 13, doi:10.1093/gbe/evab087.

16. Ratcliff, J.; Simmonds, P. Potential APOBEC-mediated RNA editing of the genomes of SARS-CoV-2 and other coronaviruses and its impact on their longer term evolution. Virology 2021, 556, 62–72, doi:10.1016/j.virol.2020.12.018.

17. Mercatelli, D.; Giorgi, F.M. Geographic and Genomic Distribution of SARS-CoV-2 Mutations. 2020, 11, doi:10.3389/fmicb.2020.01800.

18. Takada, K.; Ueda, M.T.; Watanabe, T.; Nakagawa, S. Genomic diversity of SARS-CoV-2 can be accelerated by a mutation in the nsp14 gene. 2020, 2020.2012.2023.424231, doi:10.1101/2020.12.23.424231 %J bioRxiv.

19. Jung, C.; Kmiec, D.; Koepke, L.; Zech, F.; Jacob, T.; Sparrer, K.M.J.; Kirchhoff, F. Omicron: What Makes the Latest SARS-CoV-2 Variant of Concern So Concerning? 2022, 96, e02077–02021, doi:doi:10.1128/jvi.02077-21.

20. Focosi, D. SARS-CoV-2 Spike protein convergent evolution 2021.

21. Neto, D.F.L.; Fonseca, V.; Jesus, R.; Dutra, L.H.; Portela, L.M.O.; Freitas, C.; Fillizola, E.; Soares, B.; Abreu, A.L.; Twiari, S.; et al. Molecular dynamics simulations of the SARS-CoV-2 Spike protein and variants of concern: structural evidence for convergent adaptive evolution. Journal of biomolecular structure & dynamics 2022, 1–13, doi:10.1080/07391102.2022.2097955.

22. Upadhyay, V.; Patrick, C.; Lucas, A.; Mallela, K.M.G. Convergent Evolution of Multiple Mutations Improves the Viral Fitness of SARS-CoV-2 Variants by Balancing Positive and Negative Selection. Biochemistry 2022, 61, 963–980, doi:10.1021/acs.biochem.2c00132.

23. Martin, D.P.; Weaver, S.; Tegally, H.; San, J.E.; Shank, S.D.; Wilkinson, E.; Lucaci, A.G.; Giandhari, J.; Naidoo, S.; Pillay, Y.; et al. The emergence and ongoing convergent evolution of the SARS-CoV-2 N501Y lineages. Cell 2021, 184, 5189-5200.e5187, doi:10.1016/j.cell.2021.09.003.

24. Variant report 2022-09-14. Accessed online at https://github.com/neherlab/SARS-CoV-2_variant-reports/blob/main/reports/variant_report_2022-09-14.md on November 23, 2022.

25. Resende, P.C.; Naveca, F.G.; Lins, R.D.; Dezordi, F.Z.; Ferraz, M.V.F.; Moreira, E.G.; Coêlho, D.F.; Motta, F.C.; Paixão, A.C.D.; Appolinario, L.; et al. The ongoing evolution of variants of concern and interest of SARS-CoV-2 in Brazil revealed by convergent indels in the amino (N)-terminal domain of the spike protein. Virus Evol 2021, 7, veab069, doi:10.1093/ve/veab069.

26. Cao, Y.; Jian, F.; Wang, J.; Yu, Y.; Song, W.; Yisimayi, A.; Wang, J.; An, R.; Zhang, N.; Wang, Y.; et al. Imprinted SARS-CoV-2 humoral immunity induces convergent Omicron RBD evolution. 2022, 2022.2009.2015.507787, doi:10.1101/2022.09.15.507787 %J bioRxiv.

27. Fischer, C.; Maponga, T.G.; Yadouleton, A.; Abílio, N.; Aboce, E.; Adewumi, P.; Afonso, P.; Akorli, J.; Andriamandimby, S.F.; Anga, L.; et al. Gradual emergence followed by exponential spread of the SARS-CoV-2 Omicron variant in Africa. 0, eadd8737. doi:doi:10.1126/science.add8737.

28. Focosi, D.; Maggi, F. Do We Really Need Omicron Spike-Based Updated COVID-19 Vaccines? Evidence and Pipeline. Viruses 2022, 14, doi:10.3390/v14112488.

29. Al Khalaf, R.; Bernasconi, A.; Pinoli, P.; Ceri, S. Analysis of co-occurring and mutually exclusive amino acid changes and detection of convergent and divergent evolution events in SARS-CoV-2. Computational and structural biotechnology journal 2022, 20, 4238–4250, doi:10.1016/j.csbj.2022.07.051.

30. Kim, I.-J.; Lee, Y.-h.; Khalid, M.M.; Zhang, Y.; Ott, M.; Verdin, E. SARS-CoV-2 ORF8 limits expression levels of Spike antigen. 2022, 2022.2011.2009.515752, doi:10.1101/2022.11.09.515752 %J bioRxiv.

31. Akaishi, T.; Fujiwara, K.; Ishii, T. Insertion/deletion hotspots in the Nsp2, Nsp3, S1, and ORF8 genes of SARS-related coronaviruses. BMC ecology and evolution 2022, 22, 123, doi:10.1186/s12862-022-02078-7.

32. Focosi, D.; McConnell, S.; Casadevall, A.; Cappello, E.; Valdiserra, G.; Tuccori, M. Monoclonal antibody therapies against SARS-CoV-2. The Lancet Infectious Diseases 2022, 22, 00311–00315, doi:10.1016/S1473-3099(22)00311-5.

33. Focosi, D.; Casadevall, A. A Critical Analysis of the Use of Cilgavimab plus Tixagevimab Monoclonal Antibody Cocktail (Evusheld™) for COVID19 Prophylaxis and Treatment. 2022, 14, 1999.

34. Focosi, D.; Maggi, F.; Franchini, M.; McConnell, S.; Casadevall, A. Analysis of Immune Escape Variants from Antibody-Based Therapeutics against COVID-19: A Systematic Review. International journal of molecular sciences 2022, 23, 29.

35. Copin, R.; Baum, A.; Wloga, E.; Pascal, K.E.; Giordano, S.; Fulton, B.O.; Zhou, A.; Negron, N.; Lanza, K.; Chan, N.; et al. The monoclonal antibody combination REGEN-COV protects against SARS-CoV-2 mutational escape in preclinical and human studies. Cell 2021, 184, 3949–3961, doi: 10.1016/j.cell.2021.06.002.

36. Magnus, C.L.; Hiergeist, A.; Schuster, P.; Rohrhofer, A.; Medenbach, J.; Gessner, A.; Peterhoff, D.; Schmidt, B. Targeted escape of SARS-CoV-2 in vitro from monoclonal antibody S309, the precursor of sotrovimab. Frontiers in immunology 2022, 13, 966236, doi:10.3389/fimmu.2022.966236.

37. Dong, J.; Zost, S.J.; Greaney, A.J.; Starr, T.N.; Dingens, A.S.; Chen, E.C.; Chen, R.E.; Case, J.B.; Sutton, R.E.; Gilchuk, P.; et al. Genetic and structural basis for SARS-CoV-2 variant neutralization by a two-antibody cocktail. Nat Microbiol 2021, 6, 1233–1244, doi:10.1038/s41564-021-00972-2.

38. FDA. Fact sheet for healthcare providers: emergency use authorization for Evusheld™ (tixagevimab co-packaged with cilgavimab). Accessed online at https://www.fda.gov/media/154701/download on August 13, 2022. 2021.

39. Lee, J.S.; Yun, K.W.; Jeong, H.; Kim, B.; Kim, M.J.; Park, J.H.; Shin, H.S.; Oh, H.S.; Sung, H.; Song, M.G.; et al. SARS-CoV-2 shedding dynamics and transmission in immunosuppressed patients. Virulence 2022, 13, 1242–1251, doi:10.1080/21505594.2022.2101198.

40. Andrés, C.; González-Sánchez, A.; Jiménez, M.; Márquez-Algaba, E.; Piñana, M.; Fernández-Naval, C.; Esperalba, J.; Saubi, N.; Quer, J.; Rando-Segura, A.; et al. Emergence of Delta and Omicron variants carrying resistance-associated mutations in immunocompromised patients undergoing Sotrovimab treatment with long viral excretion. Clinical microbiology and infection : the official publication of the European Society of Clinical Microbiology and Infectious Diseases 2022, doi:10.1016/j.cmi.2022.08.021.

41. Gonzalez-Reiche, A.S.; Alshammary, H.; Schaefer, S.; Patel, G.; Polanco, J.; Amoako, A.A.; Rooker, A.; Cognigni, C.; Floda, D.; van de Guchte, A.; et al. Intrahost evolution and forward transmission of a novel SARS-CoV-2 Omicron BA.1 subvariant. 2022, 2022.2005.2025.22275533, doi:10.1101/2022.05.25.22275533 %J medRxiv.

42. Lee, C.Y.; Shah, M.K.; Hoyos, D.; Solovyov, A.; Douglas, M.; Taur, Y.; Maslak, P.; Babady, N.E.; Greenbaum, B.; Kamboj, M.; et al. Prolonged SARS-CoV-2 Infection in Patients with Lymphoid Malignancies. Cancer discovery 2022, 12, 62–73, doi:10.1158/2159-8290.Cd-21-1033.

43. Sonnleitner, S.T.; Prelog, M.; Sonnleitner, S.; Hinterbichler, E.; Halbfurter, H.; Kopecky, D.B.C.; Almanzar, G.; Koblmüller, S.; Sturmbauer, C.; Feist, L.; et al. Cumulative SARS-CoV-2 mutations and corresponding changes in immunity in an immunocompromised patient indicate viral evolution within the host. Nat Commun 2022, 13, 2560, doi:10.1038/s41467-022-30163-4.

44. Fratev, F. R346K Mutation in the Mu Variant of SARS-CoV-2 Alters the Interactions with Monoclonal Antibodies from Class 2: A Free Energy Perturbation Study. Journal of chemical information and modeling 2022, 62, 627–631, doi:10.1021/acs.jcim.1c01243.

45. Li, L.; Liao, H.; Meng, Y.; Li, W.; Han, P.; Liu, K.; Wang, Q.; Li, D.; Zhang, Y.; Wang, L.; et al. Structural basis of human ACE2 higher binding affinity to currently circulating Omicron SARS-CoV-2 sub-variants BA.2 and BA.1.1. Cell 2022, 185, 2952-2960.e2910, doi:10.1016/j.cell.2022.06.023.

46. Uraki, R.; Kiso, M.; Imai, M.; Yamayoshi, S.; Ito, M.; Fujisaki, S.; Takashita, E.; Ujie, M.; Furusawa, Y.; Yasuhara, A.; et al. Therapeutic efficacy of monoclonal antibodies and antivirals against SARS-CoV-2 Omicron BA.1 in Syrian hamsters. Nat Microbiol 2022, 7, 1252–1258, doi:10.1038/s41564-022-01170-4.

47. Nutalai, R.; Zhou, D.; Tuekprakhon, A.; Ginn, H.M.; Supasa, P.; Liu, C.; Huo, J.; Mentzer, A.J.; Duyvesteyn, H.M.E.; Dijokaite-Guraliuc, A.; et al. Potent cross-reactive antibodies following Omicron breakthrough in vaccinees. Cell 2022, 185, 2116-2131.e2118, doi:10.1016/j.cell.2022.05.014.

48. Duty, J.A.; Kraus, T.; Zhou, H.; Zhang, Y.; Shaabani, N.; Yildiz, S.; Du, N.; Singh, A.; Miorin, L.; Li, D.; et al. Discovery of a SARS-CoV-2 Broadly-Acting Neutralizing Antibody with Activity against Omicron and Omicron + R346K Variants. 2022, 2022.2001.2019.476998, doi:10.1101/2022.01.19.476998 %J bioRxiv.

49. Koyama, T.; Miyakawa, K.; Tokumasu, R. S S.J.; Kudo, M.; Ryo, A. Evasion of vaccine-induced humoral immunity by emerging sub-variants of SARS-CoV-2. Future microbiology 2022, 17, 417–424, doi:10.2217/fmb-2022-0025.

50. Castelli, M.; Baj, A.; Criscuolo, E.; Ferrarese, R.; Diotti, R.; Sampaolo, M.; Novazzi, F.; Dalla Gasperina, D.; Focosi, D.; Locatelli, M.; et al. Characterization of a lineage C.36 SARS-CoV-2 isolate with reduced susceptibility to neutralization circulating in Lombardy, Italy. Viruses 2021.

51. Jian, F.; Yu, Y.; Song, W.; Yisimayi, A.; Yu, L.; Gao, Y.; Zhang, N.; Wang, Y.; Shao, F.; Hao, X.; et al. Further humoral immunity evasion of emerging SARS-CoV-2 BA.4 and BA.5 subvariants. 2022, 2022.2008.2009.503384, doi:10.1101/2022.08.09.503384 %J bioRxiv.

52. Turelli, P.; Fenwick, C.; Raclot, C.; Genet, V.; Pantaleo, G.; Trono, D. P2G3 human monoclonal antibody neutralizes SARS-CoV-2 Omicron subvariants including BA.4 and BA.5 and Bebtelovimab escape mutants. 2022, 2022.2007.2028.501852, doi:10.1101/2022.07.28.501852 %J bioRxiv.

53. Umair, M.; Ikram, A.; Salman, M.; Haider, S.A.; Badar, N.; Rehman, Z.; Ammar, M.; Rana, M.S.; Ali, Q. Genomic surveillance reveals the detection of SARS-CoV-2 delta, beta, and gamma VOCs during the third wave in Pakistan. Journal of medical virology 2022, 94, 1115–1129, doi:10.1002/jmv.27429.

54. Ortega, J.T.; Pujol, F.H.; Jastrzebska, B.; Rangel, H.R. Mutations in the SARS-CoV-2 spike protein modulate the virus affinity to the human ACE2 receptor, an in silico analysis. EXCLI journal 2021, 20, 585–600, doi:10.17179/excli2021-3471.

55. Weisblum, Y.; Schmidt, F.; Zhang, F.; DaSilva, J.; Poston, D.; Lorenzi, J.C.C.; Muecksch, F.; Rutkowska, M.; Hoffmann, H.-H.; Michailidis, E.; et al. Escape from neutralizing antibodies by SARS-CoV-2 spike protein variants. eLife 2020, 28, e61312, doi:10.7554/eLife.61312.

56. Saifi, S.; Ravi, V.; Sharma, S.; Swaminathan, A.; Chauhan, N.S.; Pandey, R. SARS-CoV-2 VOCs, Mutational diversity and clinical outcome: Are they modulating drug efficacy by altered binding strength? Genomics 2022, 114, 110466, doi:10.1016/j.ygeno.2022.110466.

57. Focosi, D.; Maggi, F. Recombination in Coronaviruses, with a focus on SARS-CoV-2. Viruses 2022, 14, 1239.

58. Roemer, C.H. R; Frohberg, N.; Sakaguchi, H.; Gueli, F.; Peacock, T. SARS-CoV-2 evolution, post-Omicron. Available online: https://virological.org/t/sars-cov-2-evolution-post-omicron/911 (accessed on November 26).

59. Lambisia, A.; Nyiro, J.; Morobe, J.; Makori, T.; Ndwiga, L.; Mburu, M.; Moraa, E.; Musyoki, J.; Murunga, N.; Bejon, P.; et al. Detection of a SARS-CoV-2 Beta-like Variant with Additional Mutations in Coastal Kenya after >1 Year of Disappearance. Available online: https://virological.org/t/detection-of-a-sars-cov-2-beta-like-variant-with-additional-mutations-in-coastal-kenya-after-1-year-of-disappearance/910 (accessed on November 26).

60. Sullivan, D.J.; Franchini, M.; Joyner, M.J.; Casadevall, A.; Focosi, D. Analysis of anti-Omicron neutralizing antibody titers in different convalescent plasma sources. Nat Comm 2022, 13, 6478, doi:10.1038/s41467-022-33864-y.

61. Sullivan, D.J.; Franchini, M.; Senefeld, J.W.; Joyner, M.J.; Casadevall, A.; Focosi, D. Plasma after both SARS-CoV-2 boosted vaccination and COVID-19 potently neutralizes BQ1.1 and XBB. 2022, 2022.2011.2025.517977, doi:10.1101/2022.11.25.517977 %J bioRxiv.

62. FDA Announces Bebtelovimab is Not Currently Authorized in Any US Region. Accessed online at https://www.fda.gov/drugs/drug-safety-and-availability/fda-announces-bebtelovimab-not-currently-authorized-any-us-region on December 1, 2022.

63. FDA Statement. January 24, 2022. Coronavirus (COVID-19) Update: FDA Limits Use of Certain Monoclonal Antibodies to Treat COVID-19 Due to the Omicron Variant. Accessed online at https://www.fda.gov/news-events/press-announcements/coronavirus-covid-19-update-fda-limits-use-certain-monoclonal-antibodies-treat-covid-19-due-omicron on February 3,.

64. FDA updates Sotrovimab emergency use authorization. March 30, 2022. Accessed online at https://www.fda.gov/drugs/drug-safety-and-availability/fda-updates-sotrovimab-emergency-use-authorization on April 26, 2022.

65. FDA releases important information about risk of COVID-19 due to certain variants not neutralized by Evusheld. Accessed online at https://www.fda.gov/drugs/drug-safety-and-availability/fda-releases-important-information-about-risk-covid-19-due-certain-variants-not-neutralized-evusheld#:~:text=Therefore%2C%20on%20June%2029%2C%202022,if%20patients%20need%20ongoing%20protection. on October 10, 2022.

66. Senefeld, J.W.; Klassen, S.A.; Ford, S.K.; Senese, K.A.; Wiggins, C.C.; Bostrom, B.C.; Thompson, M.A.; Baker, S.E.; Nicholson, W.T.; Johnson, P.W.; et al. Use of convalescent plasma in COVID-19 patients with immunosuppression. Transfusion 2021, 61, 2503–2511, doi:10.1111/trf.16525.

67. Chen, C.; Nadeau, S.; Yared, M.; Voinov, P.; Xie, N.; Roemer, C.; Stadler, T. CoV-Spectrum: analysis of globally shared SARS-CoV-2 data to identify and characterize new variants. Bioinformatics 2021, 38, 1735–1737, doi:10.1093/bioinformatics/btab856 %J Bioinformatics.

68. Hadfield, J.; Megill, C.; Bell, S.M.; Huddleston, J.; Potter, B.; Callender, C.; Sagulenko, P.; Bedford, T.; Neher, R.A. Nextstrain: real-time tracking of pathogen evolution. Bioinformatics 2018, 34, 4121–4123, doi:10.1093/bioinformatics/bty407 %J Bioinformatics.

